# Neural dynamics in ventrolateral prefrontal cortex underlie learning from feedback

**DOI:** 10.1101/2025.11.16.688684

**Authors:** Runhao Lu, Mikiko Kadohisa, Makoto Kusunoki, Daniel J. Mitchell, Alexandra Woolgar, Mark J. Buckley, John Duncan

## Abstract

Learning often depends on feedback, yet how positive and negative outcomes reorganize target representations to support later memory retrieval remains poorly understood. Accumulating evidence suggests that the ventrolateral prefrontal cortex (vlPFC) acts as a central hub linking learning and retrieval, raising the possibility that it plays a critical role in this process. Here we analysed spiking activity and local field potentials (LFPs) recorded from vlPFC while monkeys performed a multi-cycle object-learning task. During the initial learning cycle, correct and incorrect feedback elicited distinct vlPFC neural responses in both spiking and LFPs. In particular, positive feedback produced elevated theta power and enhanced phase-amplitude coupling (PAC) between theta phase and high-frequency amplitude, associated with sustained suppression of neural spiking. Incorrect feedback induced stronger beta power. Despite comparable levels of object information under both feedback conditions, decoders trained and tested within the same feedback state outperformed those tested across states, revealing feedback-dependent coding formats. State-space and cross-period generalisation analyses further showed that object representations following positive feedback were geometrically closer to and shared a common coding format with those reinstated during later retrieval, indicating that feedback reshapes neural geometry toward retrieval-compatible states. Moreover, these geometric and generalisation effects were selectively expressed on electrodes showing stronger PAC or beta power, suggesting that oscillatory coordination may regulate how feedback signals are transformed into stable target codes. Together, our results reveal how vlPFC serves as a critical bridge between learning and memory retrieval, with feedback-driven dynamics reorganizing population geometry through rhythmic coordination and bringing successful outcome states closer to future retrieval representations.

## Introduction

Learning from feedback is fundamental to adaptive, goal-directed behaviour (*1–6*). Positive and negative outcomes not only modulate learning of current goals but also shape subsequent selection and memory retrieval (*7–10*). While the neural mechanisms of reinforcement have been extensively studied (*11–13*), how feedback reorganizes cortical representations of learning targets into neural codes that support later retrieval and goal-directed behaviour remains poorly understood.

In the primate brain, feedback engages interconnected prefrontal circuits including the ventrolateral and dorsolateral prefrontal cortex (vlPFC and dlPFC), which play central roles in learning, memory, and cognitive control (*14–18*). Classical accounts proposed distinct functional specializations for these regions, with vlPFC linked to object processing and dlPFC to spatial control (*19, 20*). However, accumulating evidence indicates that vlPFC may serve a more domain-general role (*16, 17, 21, 22*), coordinating multiple cognitive operations through its widespread frontotemporal and frontoparietal connections and strong oscillatory synchronisation (*23–26*). Recent large-scale recordings across vlPFC, dlPFC, and temporal cortex (TE) during a multi-step object-learning task showed that vlPFC encoded all task-relevant features (cue, object, and location) earlier and more strongly than other regions and was the only area maintaining target information across both retrieval and feedback (*22*). These findings position vlPFC as a key integrative node bridging learning and memory retrieval.

Despite this, the role of feedback in shaping target representations within prefrontal cortex remains unresolved. During learning, positive and negative feedback elicit widespread spiking and local field potential (LFP) responses across prefrontal cortices (*4, 15–18, 27, 28*). Such responses have often been interpreted as reflecting reward valuation or outcome monitoring, but they may also transiently reconfigure cortical population geometry, creating distinct neural states that bias subsequent processing toward or away from the learning target. This idea aligns with dynamic-coding frameworks in prefrontal cortex, which propose that information is flexibly represented through time-varying rather than static activity patterns (*29–33*), implying that positive and negative feedback may establish distinct representational formats within the same cortical network.

A related but unresolved question concerns the mechanisms that stabilize these reconfigured representations. Neural oscillations have long been implicated in coordinating cortical computations during learning and plasticity (*27, 34–38*). In prefrontal cortex, theta rhythms (∼4-8 Hz) typically emerge after feedback or errors and are associated with cognitive control and updating internal representations (*27, 39–42*), whereas beta activity (∼20-30 Hz) has often been linked to transient suppression of neural ensembles that are no longer relevant (*24, 38, 43*). Phase-amplitude coupling (PAC) between theta phase and high-frequency amplitude (HFA) may provide a mechanism for integrating slow feedback-related control signals with fast ensemble activity supporting associative learning and memory updating (*44–49*). These oscillatory and cross-frequency dynamics may thus represent candidate mechanisms through which feedback transforms transient neural responses into stable representational formats that can be reinstated during later retrieval. To investigate how feedback reshapes target representations to support later retrieval, we re-analysed the large-scale neural recording dataset collected by Kadohisa et al. (*22*), in which macaque monkeys performed a multi-cycle object-learning task encompassing both learning and retrieval phases. We primarily focused on the vlPFC, as previous work showed that it was the only region maintaining target information across feedback and retrieval phases (*22*). However, we also analysed the simultaneously recorded dlPFC and TE populations to establish the circuit-level specificity of these mechanisms. We examined how positive and negative feedback differentially modulate spiking and LFP activity, and how these feedback-induced states reorganize target representations during learning and bias their geometry during subsequent retrieval. Furthermore, we asked whether feedback-related oscillatory dynamics, particularly theta, beta, and PAC, modulate the stabilisation of representations that guide future behaviour. Together, these analyses allow us to test how feedback-driven dynamics in vlPFC serve as a mechanism for reformatting neural codes to support learning and goal-directed control.

## Results

### Experimental paradigm and recordings

The present study is a novel re-analysis based on the dataset previously reported by Kadohisa et al. (*22*). Full methodological details can be found in the original report. Briefly, two monkeys performed a multi-cycle object-learning task that required discovering targets through trial-and-error feedback in the first learning cycle (Cycle 1), and subsequently retrieving the same targets based on learned cues in later cycles (Cycles 2-4) (Figure 1A). In each recording session, the animal completed a series of problems, each composed of four cycles of trials. On every trial, six objects were presented in a circular array, and the animal awaited a go cue before making a saccade to one object and maintaining fixation until visual feedback appeared. Positive feedback was indicated by a coloured square (green or yellow) that replaced the chosen object and was followed by reward delivery, whereas incorrect choices produced a red square with no reward.

**Figure 1.**
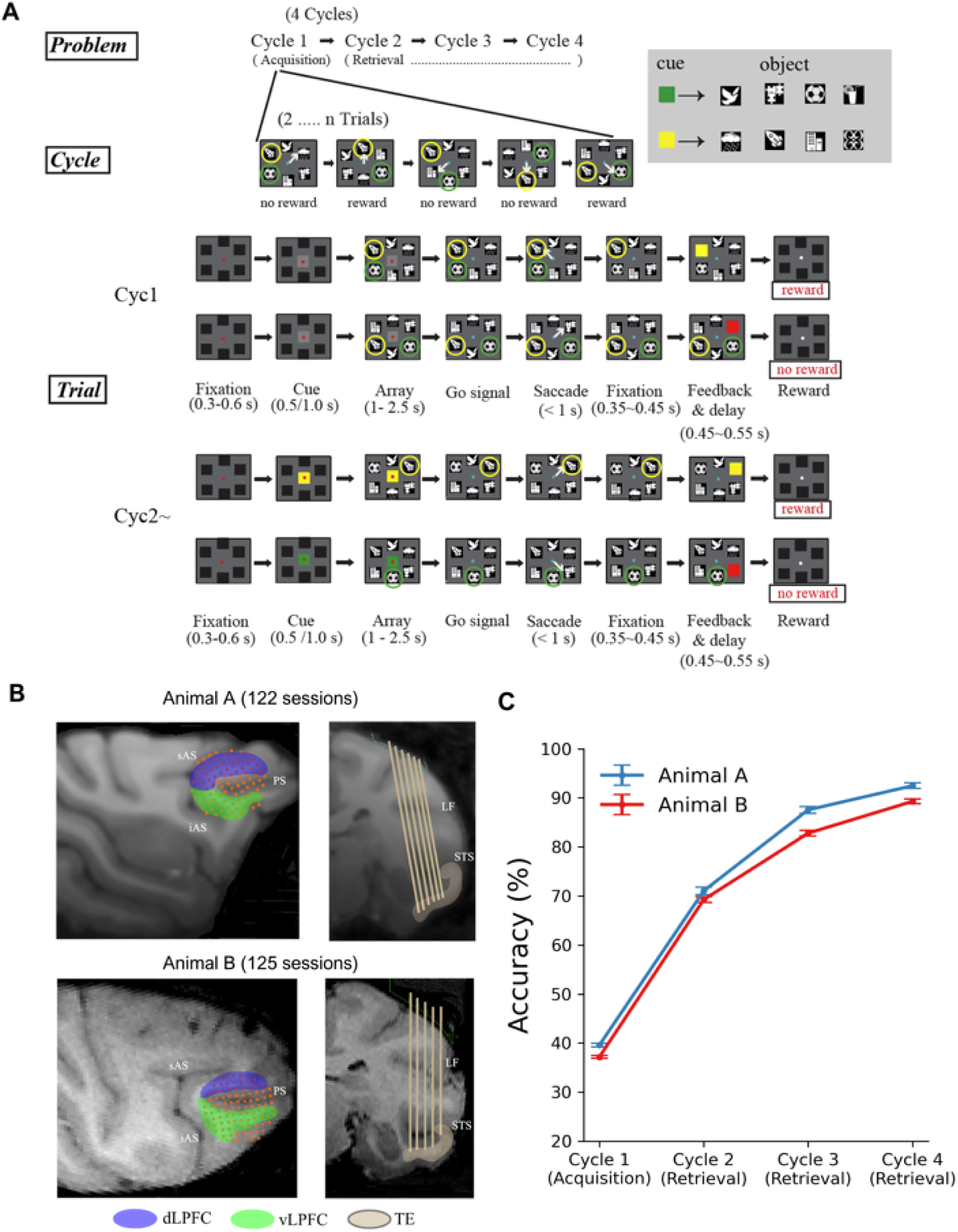
Task design and recording sites. (A) Schematic of the multi-cycle object-learning task (Adapted with permission from (*22*)). A fixed set of eight objects was used throughout the study (inset), four each associated with green and yellow cues. Each problem consisted of four cycles of trials. In the first cycle (Cycle 1, acquisition), through trial-and-error feedback, monkeys discovered two rewarded target objects, one from each four-object set (marked here with green and yellow circles, not present in real array). White arrows indicate the monkey’s saccade to a selected object. Positive feedback was indicated by a square of the appropriate cue colour (green or yellow) replacing the chosen object and followed by reward delivery, whereas incorrect selections produced a red square with no reward. In subsequent cycles (Cycles 2-4, retrieval), a coloured cue presented at fixation instructed which of the two previously rewarded objects should be selected. Each of these cycles consisted of two correct trials (plus any errors), one for each cue-target pair. After cycle 4, the problem terminated and a new pair of targets was introduced. (B) Recording locations in two monkeys (Adapted with permission from (*22*)). One semi-chronic 96-channel microelectrode array targeted the lateral prefrontal cortex, encompassing ventrolateral (vlPFC) and dorsolateral (dlPFC) prefrontal regions on the cortical convexity. The location of each electrode is shown by a red dot. Also, a separate 32-channel array was directed at the ventral surface of the temporal cortex (TE) (just a subset of electrodes is shown here on a representative coronal brain slice). sAS, superior arcuate sulcus; iAS, inferior arcuate sulcus; PS, principal sulcus. LF, lateral fissure; STS, superior temporal sulcus. (C) Learning time course across cycles. Behavioural accuracy averaged across all sessions for both animals. Cycle 1 corresponds to the trial-and-error acquisition phase. Cycles 2-4 represent the retrieval phase, during which animals successfully utilized learned cue-target associations. Error bars denote ± SEM across sessions.

Across the experiment, a fixed set of eight objects was divided into two colour-defined sets (four green and four yellow). Each problem used six of these objects, three from each colour set, arranged randomly within the array so that selections had to be based on object identity rather than spatial position. During the first learning cycle (Cycle 1), monkeys had no prior cue information and discovered the two rewarded targets (one from each colour set) through trial and error. Each target was rewarded once; after both targets had been found, the cycle ended. In subsequent retrieval cycles (Cycles 2-4), a coloured cue (green or yellow) presented at the start of each trial indicated which of the two previously rewarded objects should be selected. The cue did not reappear within the same cycle once its associated target had been chosen, so that, in the absence of error, each cycle consisted of two correct trials, one for each cue-target pair. After completion of the fourth cycle, the problem terminated, and a new pair of targets replaced the previous ones to begin a new learning episode.

Consistent with this task structure, the animals’ choice behaviour exhibited a clear transition from trial-and-error exploration to stable memory-guided selection (Figure 1C). During the initial acquisition phase (Cycle 1), overall accuracy was approximately 40%, reflecting the exploratory nature of target discovery. From Cycle 2 onwards, as the animals utilized the instruction cues, performance showed a sharp increase and stabilized at high accuracy levels (ranging from >70% to ∼90%) throughout Cycles 2, 3, and 4. Our previous report (*22*) concerned only neural activity in Cycles 2–4, the stage following trial and error learning. Here we examine consequences of positive and negative feedback in Cycle 1, when animals were first instructed on target identities.

Neural activity was recorded using semi-chronic microelectrode arrays implanted in the lateral prefrontal cortex and temporal cortices (Figure 1B). Electrodes within each array were independently movable between sessions, allowing recordings from different electrode sites across days. The original study (*22*) included recordings from the vlPFC, dlPFC, TE, and regions within the principal sulcus. Previous analyses (*22*) demonstrated that, among these regions, only the vlPFC maintained robust target object representations across both feedback and retrieval phases. Thus, the present study primarily focuses on vlPFC to examine how feedback-related neural dynamics reshape target representations that bridge learning and subsequent retrieval. Additionally, to establish the specificity of these mechanisms, we also examined feedback-driven activity and target decoding in the simultaneously recorded dlPFC and TE populations.

Specifically, one 96-channel semi-chronic array targeted extensive regions of the lateral frontal convexity, encompassing both the vlPFC (corresponding to areas 45a, 12l, and 12r) and the dlPFC (areas 46, 9, and 8), as confirmed by MRI and post-mortem histology. Also, a separate 32-channel array was directed at the ventral surface of the temporal lobe to sample from area TE (including subdivisions TEad, TEav, TEpd, and TEpv). Across 122 sessions from animal A and 125 sessions from animal B, we recorded 1252 neurons from vlPFC (933 from animal A; 319 from animal B), 1545 neurons from dlPFC (1,218 from animal A; 327 from animal B), and 372 neurons from TE (61 from animal A; 311 from animal B). After preprocessing and exclusion of noisy or unstable channels, LFP analyses were conducted on 946 vlPFC electrode locations (729 from animal A; 217 from animal B), 1200 dlPFC electrode locations (980 from animal A; 220 from animal B), and 126 TE electrode locations (57 from animal A; 69 from animal B), hereafter simply referred to as “electrodes”.

### Distinct spiking and LFP responses to positive and negative feedback in vlPFC

We first examined how positive and negative feedback modulated neural activity during the feedback period of the initial learning cycle (Cycle 1). As shown in Figure 2, feedback onset produced pronounced changes in both spiking and LFP signals in vlPFC. Population firing rates increased transiently after both correct and incorrect feedback, peaking around 140 ms after feedback onset and returning to around baseline levels by about 250ms. Thereafter, responses diverged: activity following incorrect feedback remained around baseline, whereas activity after correct feedback continued to decline and remained significantly lower during the late feedback window (250-400 ms; Figure 2A). Because prior-trial rewards can bias neural activity in prefrontal circuits (*50*), we performed a single-trial-back analysis to ensure our results were not confounded by reward history (Figure S1). We categorized neural responses in Cycle 1 according to both the current and the preceding (N-1) trial outcomes, and found that the late suppression of firing rates for current trial correct feedback remained robust during the 250–450 ms window, regardless of the prior trial’s outcome. These results indicate that the observed neural states in vlPFC are primarily driven by the evaluation of current feedback rather than lingering signals from previous trials.

**Figure 2.**
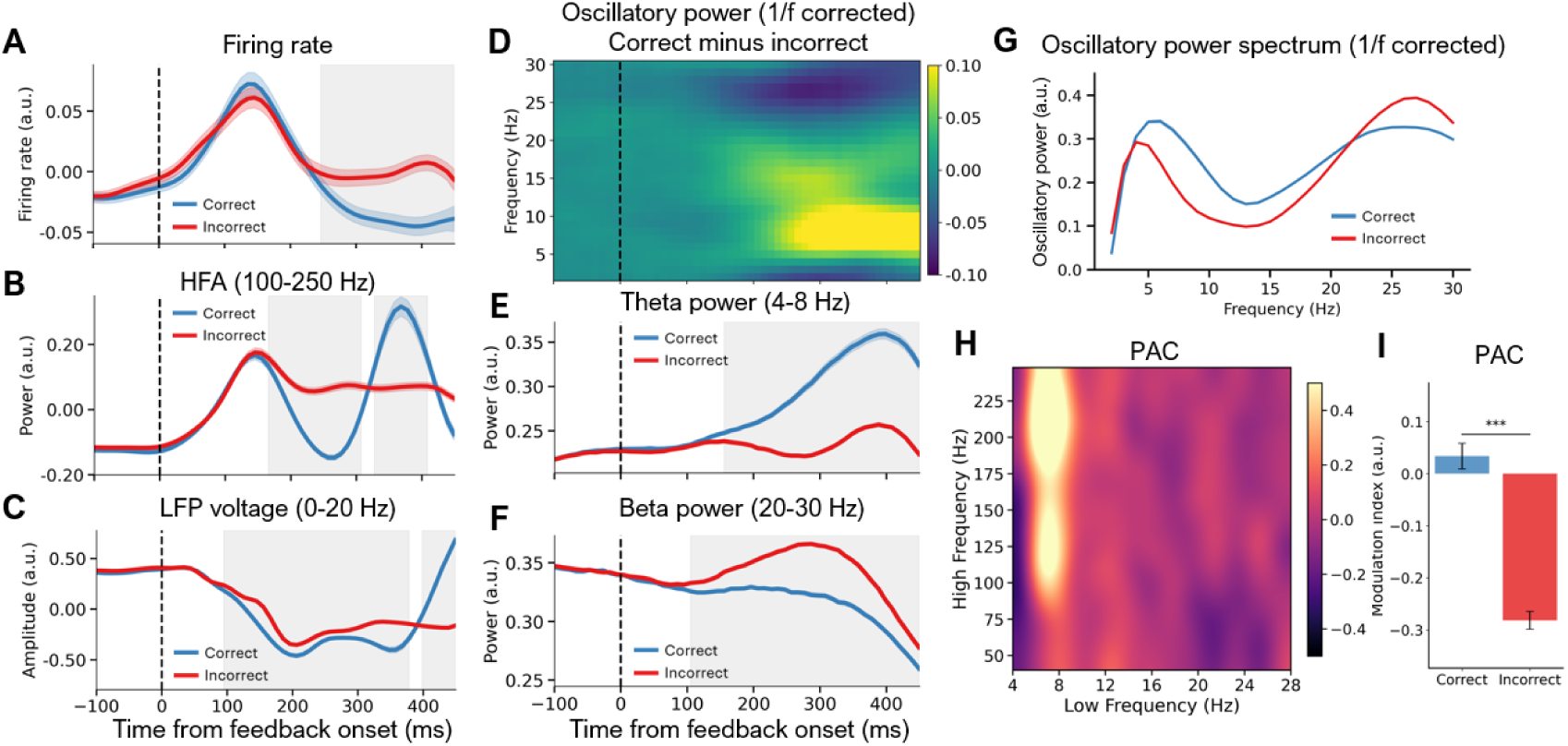
Spiking and LFP responses to positive and negative feedback in vlPFC. (A) Population firing rates averaged across vlPFC neurons for correct and incorrect feedback. (B) High-frequency activity (HFA, 100-250 Hz) averaged across vlPFC electrodes for correct and incorrect feedback. (C) Event-related potentials (ERPs) averaged across vlPFC electrodes for correct and incorrect feedback. (D) Time-frequency decomposition results of oscillatory power (correct minus incorrect feedback conditions) after removing the aperiodic 1/f component using FOOOF. (E-F) Time courses of theta (4-8 Hz) and beta (20-30 Hz) power for correct and incorrect feedback. (G) Aperiodic-corrected power spectra computed using the 200-450 ms post-feedback window. (H) Cross-frequency phase-amplitude coupling (PAC) between low (4-28 Hz) and high (40-250 Hz) frequency ranges, shown as difference between positive and negative feedback. (I) Theta-HFA PAC (4-8 Hz phase, 100-250 Hz amplitude) during 200-450 ms window for correct and incorrect feedback. All time 0 values correspond to feedback onset in Cycle 1 (the learning cycle). Gray shading regions indicate time windows with significant differences between correct and incorrect trials (corrected *p* < 0.05, cluster-based permutation test). Significance was assessed by constructing empirical null distributions from permutation samples while accounting for serial-position effects (see Methods). Error shading or error bars denote ± SEM across neurons or electrodes. *** *p* < 0.001.

In part, high-frequency activity (HFA, 100-250 Hz) paralleled the spiking response. Following feedback onset, HFA rose sharply for both correct and incorrect trials, peaking at ∼140 ms and aligning well with the population firing-rate peak (Figure 2A-B). After this initial rise, HFA quickly plateaued in incorrect trials, whereas after correct feedback it displayed a rhythmic modulation suggestive of coupling with slower oscillations in the theta range.

LFP voltage (i.e., event-related potentials, ERP) also differentiated feedback valence (Figure 2C). Both correct and incorrect feedback elicited a clear negative-going deflection beginning shortly after feedback onset. This deflection was significantly larger for correct trials and extended over ∼100-380 ms. Around 400 ms, correct-feedback trials showed a pronounced polarity reversal, producing a late positive component that was absent following incorrect feedback.

To characterize the oscillatory activity of the LFP more precisely, we applied time-resolved fitting oscillations & one over f (FOOOF) analysis (*51, 52*) to separate periodic oscillations from the aperiodic 1/*f* background (Figure 2D). After removing the aperiodic component, we observed two distinct oscillatory differences during the feedback period: a relative increase in theta power and a relative decrease in beta power for correct compared with incorrect feedback. Time-resolved analyses of each condition separately showed that theta power increased markedly after correct feedback but changed little after incorrect feedback, yielding a significant difference between conditions from ∼160 to 450 ms (Figure 2E). In contrast, beta power decreased after correct feedback but rose transiently after incorrect feedback, peaking around 290 ms before declining, with significant differences persisting throughout ∼100-450 ms (Figure 2F).

To confirm the frequency specificity of these effects, we examined the aperiodic-corrected power spectra averaged over the 200-450 ms window of the feedback period (Figure 2G). The spectra revealed clear peaks confined to the theta and beta ranges, with no evidence for an alpha-band peak, indicating that feedback-related oscillations in vlPFC were dominated by these two frequencies. Subsequent analyses therefore focused on theta and beta activity.

We next examined the strength of phase-amplitude coupling during the 200-450 ms feedback window across a low-frequency range of 4-28 Hz and a high-frequency range of 40-250 Hz using modulation index (MI) (*53*). The difference map between positive and negative feedback in Figure 2H revealed a selective increase in the theta-HFA sector, with no reliable effects in other cross-frequency combinations. To quantify this effect, we computed PAC using theta phases (4-8 Hz) and HFA (100-250 Hz). We found that PAC was significantly higher for correct than incorrect feedback, consistent with the oscillatory modulation of HFA on correct trials (Figure 2B), and confirming enhanced theta-HFA coordination associated with successful outcomes (Figure 2I). To verify the robustness of this broadband effect, we further split the high-frequency amplitude into lower (100-150 Hz) and upper (175-250 Hz) gamma bands. We found that the feedback-dependent enhancement of PAC remained highly significant across both sub-bands, with the upper range (175-250 Hz) exhibiting a particularly pronounced difference between feedback conditions (Figure S2).

To rule out the possibility that these univariate differences were driven by the unequal number of correct and incorrect trials inherent to the trial-and-error nature of Cycle 1, we performed a subsampled control analysis on both spiking and LFP signals. By strictly matching trial counts (1:1) across feedback conditions and object identities via random subsampling, we confirmed that these distinct spiking and LFP voltage dynamics remained highly robust (Figure S3).

As another control analysis, we further investigated whether the feedback-induced theta and beta oscillations reflect coordinated or independent processes by analysing their spatial correlation and cross-frequency coupling. We observed a robust negative correlation between theta and beta power across vlPFC electrodes (Figure S4A), alongside a complete absence of significant theta-beta phase-amplitude coupling (Figure S4B). These results demonstrate that rather than acting as a single nested complex, theta and beta rhythms operate as functionally distinct and potentially competitive processes in response to different feedback.

Together, these results show that positive and negative feedback engage distinct spiking and LFP responses in vlPFC. Correct feedback was accompanied by enhanced theta, HFA, and PAC activity, signals typically associated with learning and representational updating (*27, 42*), alongside a sustained suppression of spiking. Incorrect feedback elicited stronger beta activity consistent with inhibitory control (*24, 43*). These complementary feedback-related patterns suggest coordinated changes in local firing and oscillatory dynamics that may set the stage for the reorganisation of object representations in subsequent learning and retrieval cycles.

### Spiking and LFP responses to positive and negative feedback in dlPFC and TE

To determine whether these feedback-induced dynamics were specific to the vlPFC or reflected broader network processes, we repeated these univariate and time-frequency analyses for the simultaneously recorded dlPFC and TE populations (Figures S5 and S6). We found that basic feedback-valence signals were widely seen across the frontotemporal network. Consistent with observations in vlPFC, both dlPFC and TE exhibited higher firing rates following incorrect compared to correct feedback (Figures S5A and S6A). In dlPFC, this valence difference emerged as early as ∼100 ms and persisted through the feedback window, whereas in TE, the significant difference was more transient, occurring between approximately 150-300 ms. Low-frequency oscillatory dynamics were also highly consistent across the three regions, with all areas exhibiting distinct theta and beta peaks following feedback onset. In both dlPFC and TE, theta power was significantly higher for correct trials, while beta power was selectively elevated after incorrect feedback (Figures S5D-G and S6D-G). Regarding HFA, dlPFC followed a temporal profile similar to vlPFC, with lower activity for correct trials from 100-300 ms followed by a late-window increase (300-450 ms; Figure S5B). In contrast, TE HFA showed a different pattern, where activity for correct trials remained elevated from ∼220 ms onwards (Figure S6B). Regional differences were also evident in LFP voltages. In dlPFC, LFP voltages were more positive for correct trials between around 160-300 ms before reversing late in the window (Figure S5C), whereas TE LFP voltages were generally more positive for correct trials throughout the period (Figure S6C). Finally, we examined cross-frequency coordination. Neither dlPFC nor TE exhibited strong Theta-HFA PAC, as evidenced by negative normalized MI values in both regions (Figures S5H-I and S6H-I). However, while TE showed no significant feedback-related difference in PAC, dlPFC exhibited a significant relative enhancement of PAC for correct compared to incorrect feedback, mirroring the directional effect seen in vlPFC. These comparisons indicate that much of the feedback-related neural dynamics are repeated across the frontotemporal network, particularly in dlPFC, though some signals (e.g., LFP voltages and the strength of PAC) exhibit notable regional variations.

### Functional heterogeneity of feedback responses across neurons and electrodes

To further dissect the population dynamics, we systematically characterized the functional heterogeneity across both individual neurons and LFP channels during the post-feedback period, as previous studies have highlighted heterogeneous or opposite response directions in prefrontal circuits processing feedback (*54, 55*). Specifically, to quantify the reliable differentiation between correct and incorrect outcomes during the 200–450 ms post-feedback window, we performed independent t-test across trials for each neuron and electrode for firing rate, LFP voltage, oscillatory power, and HFA. For PAC, because single-trial estimation inherently suffers from low signal-to-noise ratios, we computed the coupling strength using all trials within each condition per electrode (identical to the method used for the analysis in Figure 2I), and evaluated the significance of the condition difference against an empirical null distribution generated via trial-label permutation (see Methods).

In the vlPFC, a substantial proportion of neurons significantly differentiated between feedback states. Specifically, 28.5% of neurons exhibited significantly higher firing rates for incorrect feedback, while 18.5% preferred correct feedback (Figure S7A). This larger proportion of incorrect-preferring neurons aligns with the population-level suppression after correct feedback reported in our main results (Figure 2A). For LFP features, the proportions of electrodes showing significant preferences for either correct or incorrect feedback were more balanced for LFP voltage (42.6% vs. 35.3%) and HFA (33.6% vs. 35.8%) (Figure S7B, C). This relatively balanced distribution likely reflects the complex temporal dynamics of these signals; since both LFP voltages and HFA exhibit prominent direction reversals within the 200–450 ms window (see Figure 2B, C), these two sub-populations may effectively capture distinct early and late temporal phases of the feedback response. In contrast, frequency-specific oscillatory power exhibited the most consistent and uni-directional responses. Theta power was significantly enhanced for correct feedback in a vast majority of responsive electrodes (40.2% vs. 1.9%; Figure S8A), while beta power was overwhelmingly greater following incorrect feedback (33.0% vs. 2.2%; Figure S8B). For theta-HFA PAC, 22.9% of electrodes significantly preferred correct feedback, while 8.6% preferred incorrect feedback (Figure S8C).

Similar patterns of heterogeneity were observed in the dlPFC and TE. For spiking activity, both regions showed a higher proportion of neurons preferring incorrect feedback (dlPFC: 24.6% vs. 11.5%; TE: 24.7% vs. 14.5%; Figure S7A). Consistently, LFP voltage and HFA in these regions also exhibited mixed response directions (Figure S7B, C), while oscillatory power maintained the uni-directional dominance for theta and beta bands observed in the vlPFC (Figure S8A, B). Theta-HFA PAC exhibited distinct regional patterns of feedback selectivity: a slightly higher proportion of dlPFC electrodes showed stronger coupling for correct feedback (14.6% vs. 12.9%), whereas TE electrodes predominantly preferred correct feedback (14.3% vs. 4.8%; Figure S8C).

Together, these findings demonstrate that while stable group-level effects are evident in our main results, individual neurons and electrodes exhibit marked functional heterogeneity. Specifically, many individual sites show differential responses to correct versus incorrect feedback, with certain signals (such as theta and beta power) displaying a strong dominant response direction. Nevertheless, regardless of the specific direction or preference, a substantial proportion of neurons and electrodes reliably distinguish between the two feedback states. This widespread, single-site selectivity provides the necessary neural substrate for the distinct, feedback-dependent representational formats and geometric reorganisation observed at the population level.

### Feedback-dependent object representations in vlPFC

We next examined how feedback valence influenced representations of the target object in vlPFC (Figure 3). Object information (the identity of each object within each colour set) was decoded separately from spiking activity and LFP voltage during the feedback period of Cycle 1. For both signals, target decoding accuracy was substantially above chance under both correct and incorrect feedback conditions (Figure 3A, B, left), suggesting that object-specific information was represented before and after feedback regardless of outcome. However, despite similar overall decoding performance, representational formats differed between the two feedback states. Classifiers trained and tested within the same feedback condition (within-feedback decoding) yielded higher accuracy than classifiers trained in one condition and tested in the other (between-feedback decoding; Figure 3A, B, right), although decoding accuracies for both within- and between-feedback classifiers remained significantly above chance. This effect was consistent across spikes and LFPs and was seen from ∼160 to 400 ms after feedback onset. These results suggest that feedback modulates the underlying population representation of target objects, such that positive and negative outcomes bias the representational format in different ways.

**Figure 3.**
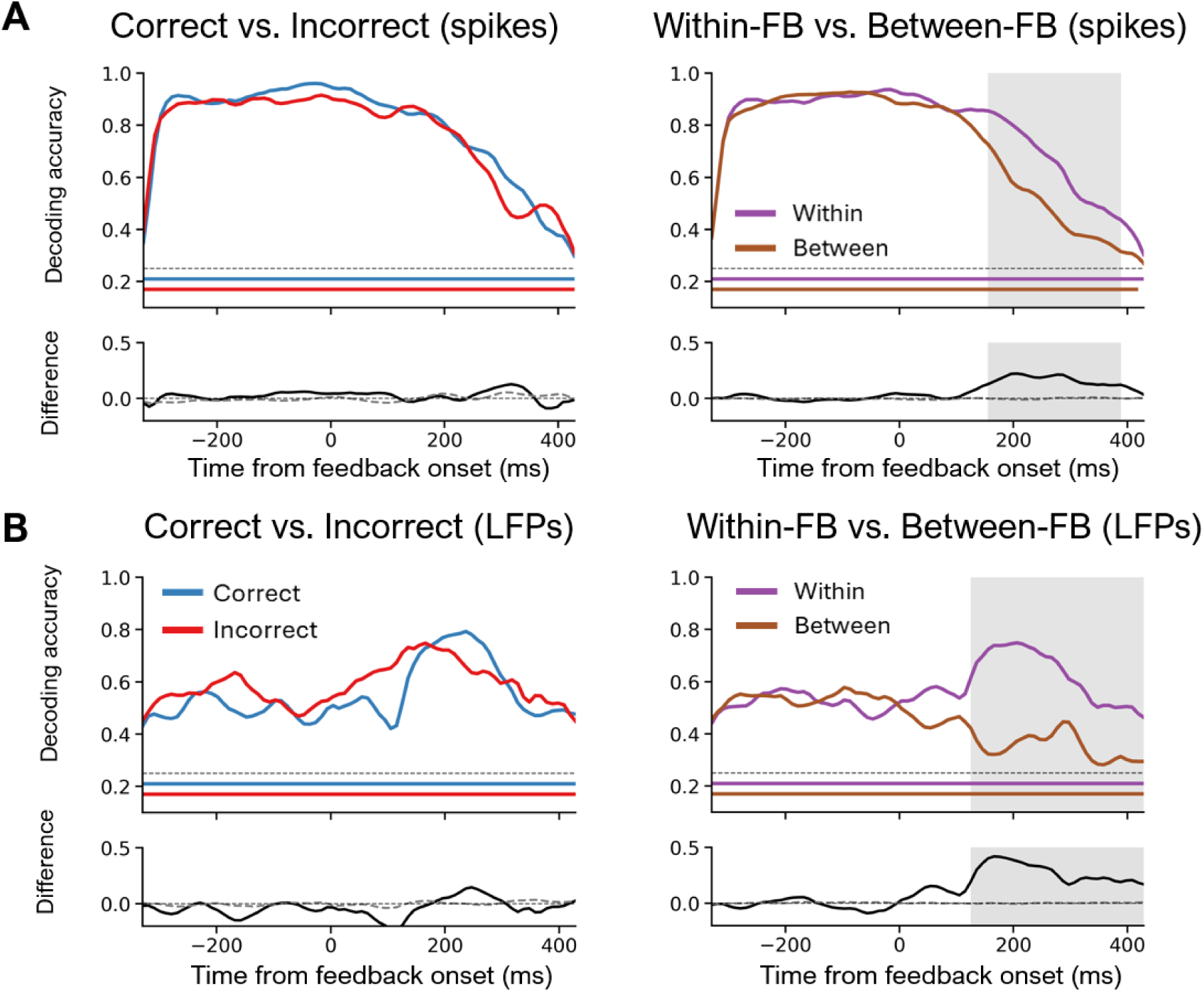
Feedback-dependent object representations in vlPFC. (A) Firing-rate-based decoding of target objects during the feedback period in Cycle 1. Left, decoding accuracy for correct (blue) and incorrect (red) feedback trials, with the lower panel showing their difference. Right, decoding accuracy when classifiers were trained and tested within the same feedback condition (within-FB) or across conditions (between-FB), with the lower panel showing within-minus-between differences. (B) Same analyses as in (A) but based on LFP voltage signals. All analyses were aligned to feedback onset (time 0). Horizontal dotted line indicates theoretical chance level (0.25). Coloured horizontal bars indicate time windows in which decoding accuracy was significantly above chance (corrected *p* < 0.05, cluster-based permutation test). Shaded regions denote time windows with significant differences between conditions (corrected *p* < 0.05, cluster-based permutation test). In the difference panels, the grey dashed line shows the permutation-derived baseline (chance-level). Tests against chance were two-tailed, whereas tests for condition differences were one-tailed.

Together, these results indicate that although target information persists in vlPFC following both positive and negative feedback, it is embedded within distinct representational states shaped by the preceding feedback. These state-dependent formats suggest that feedback reorganizes object codes in ways that constrain their cross-state generalisation, potentially biasing how information is stabilized or accessed in memory.

### Reduced sustained and state-dependent target representations in dlPFC and TE

We next examined whether this state-dependent representational formatting (within-feedback > between-feedback generalisation) was a general property of the broader network. In dlPFC, spike-based target decoding was generally weak and fell toward baseline during the post-feedback period, lacking consistent state-dependency (Figure S5J-K). While LFP signals in dlPFC showed slightly more sustained information and brief periods of state-dependent coding (Figure S5L-M), it did not exhibit the robust, persistent representational formats observed in the vlPFC. In TE, spike-based target decoding was highly accurate before feedback (reflecting strong visual responses to the foveated target) but dropped rapidly to chance levels around 200 ms after feedback onset, indicating an inability to stably maintain target information once the physical stimulus was removed (Figure S6J). Consequently, the state-dependent representational formatting seen in vlPFC was entirely absent in TE (Figure S6K). While LFP-based decoding in TE was significantly above chance after feedback onset, it was substantially weaker than in vlPFC and lacked any significant state dependency (Figure S6L-M). These results reveal that stable post-feedback target representations are unique to the vlPFC. Because object coding in dlPFC and TE largely decayed to baseline levels during the late feedback period, we focused our subsequent state-space analyses exclusively on the vlPFC population.

### Positive feedback reshapes object representations in vlPFC toward retrieval states

To test whether feedback reorganizes target representations to support later memory retrieval, we compared population activity patterns during feedback (Cycle 1) with those during the retrieval phase of later cycles (Cycles 2-4, array period; Figure 4). Neural activity during the retrieval period (−200 to 100 ms from array onset, Cycles 2-4) was used to define an object space by principal component analysis (PCA) applied to either spiking rates (Figure 4A) or LFP voltages (Figure 4C). Object representations from three time periods—Cycle 1 feedback after correct trials (200-450 ms from feedback onset; C1FB_corr), Cycle 1 feedback after incorrect trials (200-450 ms from feedback onset; C1FB_incorr), and Cycles 2-4 array presentations (−200 to 100 ms from array onset; C234A)—were then projected into this retrieval-based object space (i.e., C234A space). Because the eight objects were divided into two colour-defined cue sets, the analysis was performed separately for each set. The amount of variance captured by the three principal components (PCs) and the subspace principal angles between the two object sets are detailed in Supplementary Table 1.

**Figure 4.**
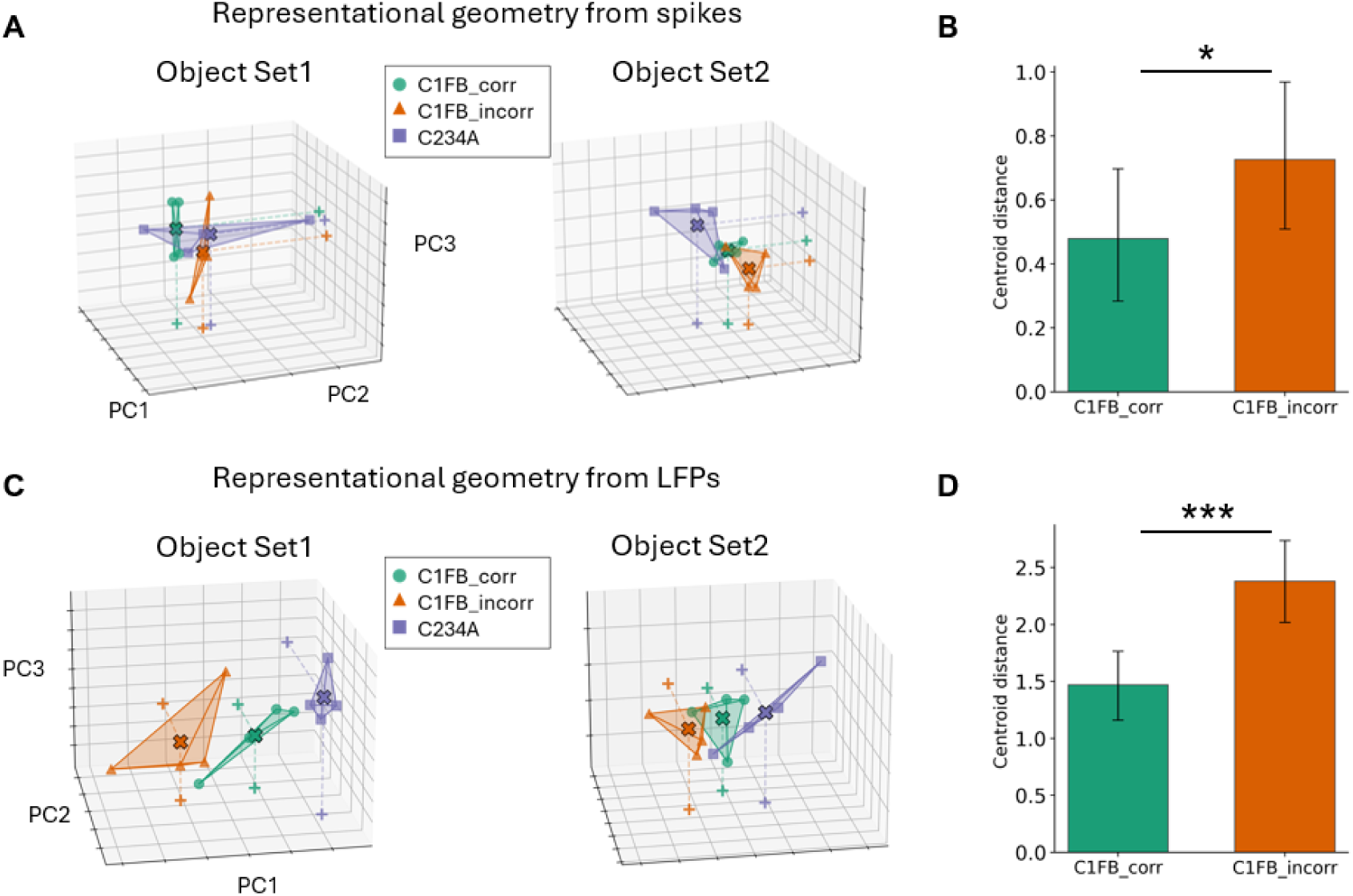
Positive feedback reshapes object representations toward retrieval states. (A, C) Object spaces constructed by principal component analysis (PCA) from population activity during the retrieval period of later cycles (Cycles 2-4 peri-array period, −200 to 100 ms from array onset). Projections are shown separately for Object Set 1 and Set 2 based on (A) spiking activity and (C) LFP voltage. For each set, object representations from three periods [Cycle 1 feedback after correct trials (C1FB_corr), Cycle 1 feedback after incorrect trials (C1FB_incorr), and Cycles 2-4 peri-array presentations (C234A)] were projected into this retrieval-based space. In each 3-D plot, different colours correspond to activity from the three task periods, and each colour’s four points represent the four objects within that object set. The shaded volumes represent the 3-D convex hulls encompassing these four objects, illustrating the representational space occupied by each task period. For each task period, the spatial centroid of the four-object constellation is marked with a bold ‘X’. To aid 3-D depth perception and clarify their relative positions, dashed lines project these centroids onto the bottom and back coordinate planes, with the projected landing points indicated by ‘+’ symbols. (B, D) Mean centroid distances between C234A and C1FB_corr or C1FB_incorr representations for (B) spike-based and (D) LFP-based object spaces. Error bars denote ± 95% confidence interval across bootstrapped samples. Statistical significance was assessed using bootstrapping analyses to generate empirical null distributions (see Methods). * *p* < 0.05, *** *p* < 0.001.

In this object space, the geometry of representations differed according to feedback valence. Across both spiking rates (Figure 4A) and LFP voltages (Figure 4C), C234A object representations were consistently closer to those from C1FB_corr than to those from C1FB_incorr. This pattern suggests that positive feedback reshaped target codes in vlPFC so that their geometry became more aligned with the states later reinstated during memory retrieval.

To quantify this relationship, we computed centroid distances between population activity patterns from each period based on bootstrapping procedures that estimated empirical distributions (see Methods). For each object set, we first averaged activity coordinates across the four objects within a period and then calculated the Euclidean distance between centroids across periods, averaging across the two sets. Both spike-based (Figure 4B) and LFP-based analyses (Figure 4D) showed significantly shorter distances between C234A and C1FB_corr than between C234A and C1FB_incorr (*M* = −0.25, 95% CI[Δ] = [−0.48, −0.01] for spike-based analysis; *M* = −0.91, 95% CI[Δ] = [−1.29, −0.52] for LFP-based analysis). These findings indicate that positive feedback drives representational convergence toward retrieval states, whereas negative feedback shifts object codes farther away, consistent with a feedback-dependent reformatting of target representations in vlPFC.

### Beta power and PAC modulate feedback-driven reorganisation of object space in vlPFC

Given that theta and theta-HFA PAC increased after correct feedback, whereas beta power was stronger after incorrect feedback (Figure 2), we next examined whether these oscillatory features modulated the feedback-dependent reshaping of object representations observed in Figure 4.

Because PAC was most enhanced following correct feedback, we first ranked all vlPFC electrodes by their PAC magnitude during 200-450 ms after positive feedback. The top one-third were defined as PAC+ electrodes and the bottom one-third as PAC− electrodes. When object spaces were constructed using only PAC+ electrodes, the strong convergence of C1FB_corr toward retrieval representations (C234A) was reproduced (*M* = −0.84, 95% CI[Δ] = [−1.17, −0.50]; Figure 5A). In contrast, object spaces derived from PAC− electrodes showed no significant difference between C1FB_corr and C1FB_incorr (*M* = 0.09, 95% CI[Δ] = [−0.20, 0.37]; Figure 5B), indicating that enhanced theta-HFA PAC is selectively associated with representational convergence toward retrieval states.

**Figure 5.**
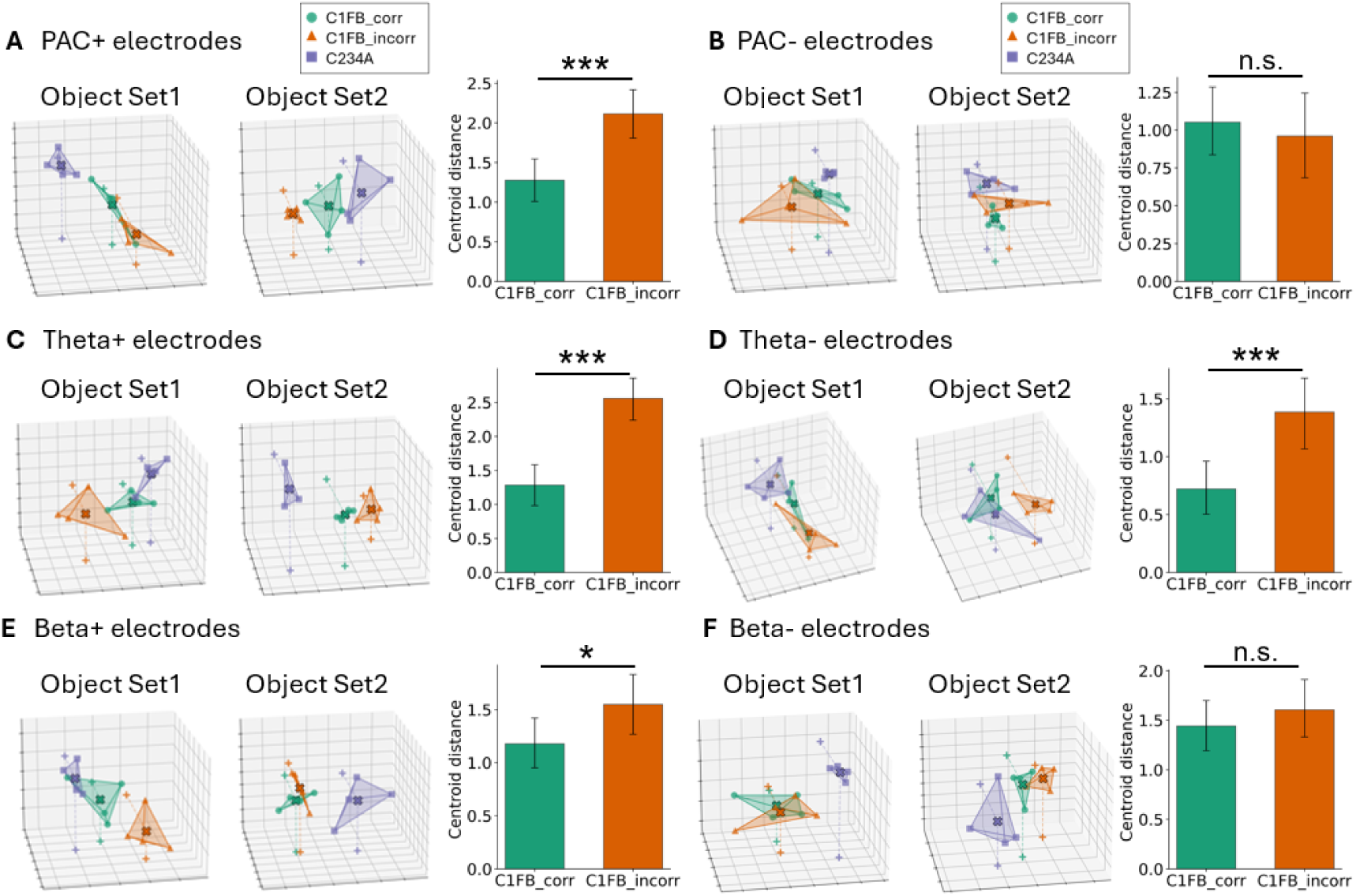
Oscillatory and cross-frequency dynamics modulate feedback-dependent reorganisation of object representations. (A, B) Object spaces constructed from vlPFC electrodes showing the strongest (PAC+) or weakest (PAC−) theta-HFA phase-amplitude coupling during 200-450 ms after correct feedback. (C, D) Object spaces constructed from electrodes with the highest (Theta+) or lowest (Theta-) theta power during 200-450 ms after correct feedback. (E, F) Object spaces constructed from electrodes with the highest (Beta+) or lowest (Beta-) beta power during 200-450 ms after incorrect feedback. Each representation geometry plot shows PCA-based projections of object representations from Cycle 1 feedback after correct (C1FB_corr) and incorrect (C1FB_incorr) trials and from retrieval periods in later cycles (C234A). Different colours correspond to activity from different task periods. Within each colour, the four points represent the four objects belonging to a given object set. The shaded volumes represent the 3-D convex hulls encompassing these four objects, illustrating the representational space occupied by each task period. For each task period, the spatial centroid of the four-object constellation is marked with a bold ‘X’. To aid 3-D depth perception and clarify their relative positions, dashed lines project these centroids onto the bottom and back coordinate planes, with the projected landing points indicated by ‘+’ symbols. Bar plots display corresponding centroid distances between feedback- and retrieval-period representations. Error bars denote ± 95% confidence interval across bootstrapped samples. Statistical significance was assessed using bootstrapping procedures to generate empirical null distributions. * *p* < 0.05, *** *p* < 0.001.

We next repeated this analysis based on theta power. Electrodes were sorted by theta power during 200-450 ms after correct feedback, with the highest one-third designated Theta+ and the lowest one-third Theta-. Because animal A exhibited a subset of electrodes with unusually strong theta activity (see Figure S9), these electrodes were excluded from ranking to ensure reliability. As shown in Figure 5C and 5D, object spaces constructed from Theta+ and Theta− electrodes both showed the convergence effect (*M* = −1.28, 95% CI[Δ] = [−1.61, −0.96] for Theta+ electrodes; *M* = −0.66, 95% CI[Δ] = [−1.00, −0.30] for Theta− electrodes), suggesting that theta power itself may not directly govern this reorganisation.

Finally, we tested whether beta power modulated the same phenomenon. Electrodes were ranked by beta power following negative feedback, and the upper and lower one-third defined as Beta+ and Beta− electrodes, respectively. Object spaces based on Beta+ electrodes alone reproduced the convergence effect (*M* = −0.37, 95% CI[Δ] = [−0.67, −0.07]), whereas those based on Beta-electrodes did not (*M* = −0.16, 95% CI[Δ] = [−0.46, 0.13]) (Figure 5E, F), indicating that stronger beta activity also influenced how feedback reshaped representational geometry.

To rule out the possibility that these distinct oscillatory features merely reflect redundant measurements from a single or highly overlapping neural population, we quantified the physical overlap among the PAC+, Theta+, and Beta+ electrode subsets (Figure S10). We found that the pairwise overlap between these subsets was relatively low, ranging from 15.5% to 22.3%. Moreover, only 26 electrodes (4.2%) were shared across all three subsets. Anatomical mapping of the most responsive electrodes (Figure S11) further revealed that these functional signatures are broadly distributed and spatially intermingled across the vlPFC.

It should be noted that, while HFA also responded robustly to feedback, we did not group electrodes by time-averaged HFA because HFA primarily reflects local spiking rather than coordinating rhythms. Moreover, time-averaging would obscure the temporally precise, rhythmic modulation of HFA observed during correct feedback (Figure 2B). Therefore, we used PAC+ vs. PAC− electrodes to explicitly quantify the phase-dependent nesting of these high-frequency activity within slower oscillations.

Together, these results suggest that oscillatory and cross-frequency dynamics in vlPFC contribute to how feedback shapes target representations. Electrodes showing stronger PAC after correct feedback, as well as those showing stronger beta power after incorrect feedback, both exhibited clearer convergence of feedback-period representations toward retrieval states. These findings indicate that multiple oscillatory processes within vlPFC may jointly support the reorganisation of object codes that underlies feedback-guided learning and subsequent memory retrieval.

### Cross-period generalisation confirms feedback-driven representational alignment modulated by oscillatory dynamics

To further quantify the observation that positive feedback aligns target representations toward future retrieval states, we performed a cross-period generalisation analysis. This approach allowed us to directly test whether the neural codes utilized during later memory retrieval (Cycles 2-4) could reliably decode the object representations formed during the initial learning feedback period (Cycle 1). To enhance decoding sensitivity, here we applied a heterogeneous reservoir computing (HeteroRC) approach (*56*) to automatically capture rich spatiotemporal features from LFP data, followed by a standard linear classifier for object decoding. Classifiers were trained on peri-array neural activity during the retrieval phase (C234A; −200 to 100 ms relative to array onset) and tested on the feedback period of the learning phase (C1FB; 200 to 450 ms relative to feedback onset) for both correct and incorrect trials. These cross-period generalisation analyses were conducted separately for the two object sets and subsequently averaged. Crucially, before decoding, we removed the centroid shift at each time point by subtracting the overall mean amplitude across all four objects within each trial. This step ensures that any observed cross-generalisation is driven by object-specific representational structures rather than being confounds of a global, non-specific signal shift following different feedback types.

Consistent with our state-space geometric analyses, cross-period generalisation was highly dependent on feedback valence (Figure 6A). Classifiers trained on the retrieval phase successfully generalized to the correct feedback state (C1FB_corr), revealing above-chance decoding accuracy. Conversely, when the same retrieval-trained classifiers were tested on the incorrect feedback state (C1FB_incorr), decoding accuracy frequently dropped below chance. This below-chance generalisation implies that negative feedback actively pushes the population representation away from the retrieval-compatible state. The direct comparison between these two conditions yielded a robust and statistically significant difference map (Figure 6A, right panel), indicating that positive feedback establishes a representational format functionally aligned with future retrieval, whereas negative feedback repels it.

**Figure 6.**
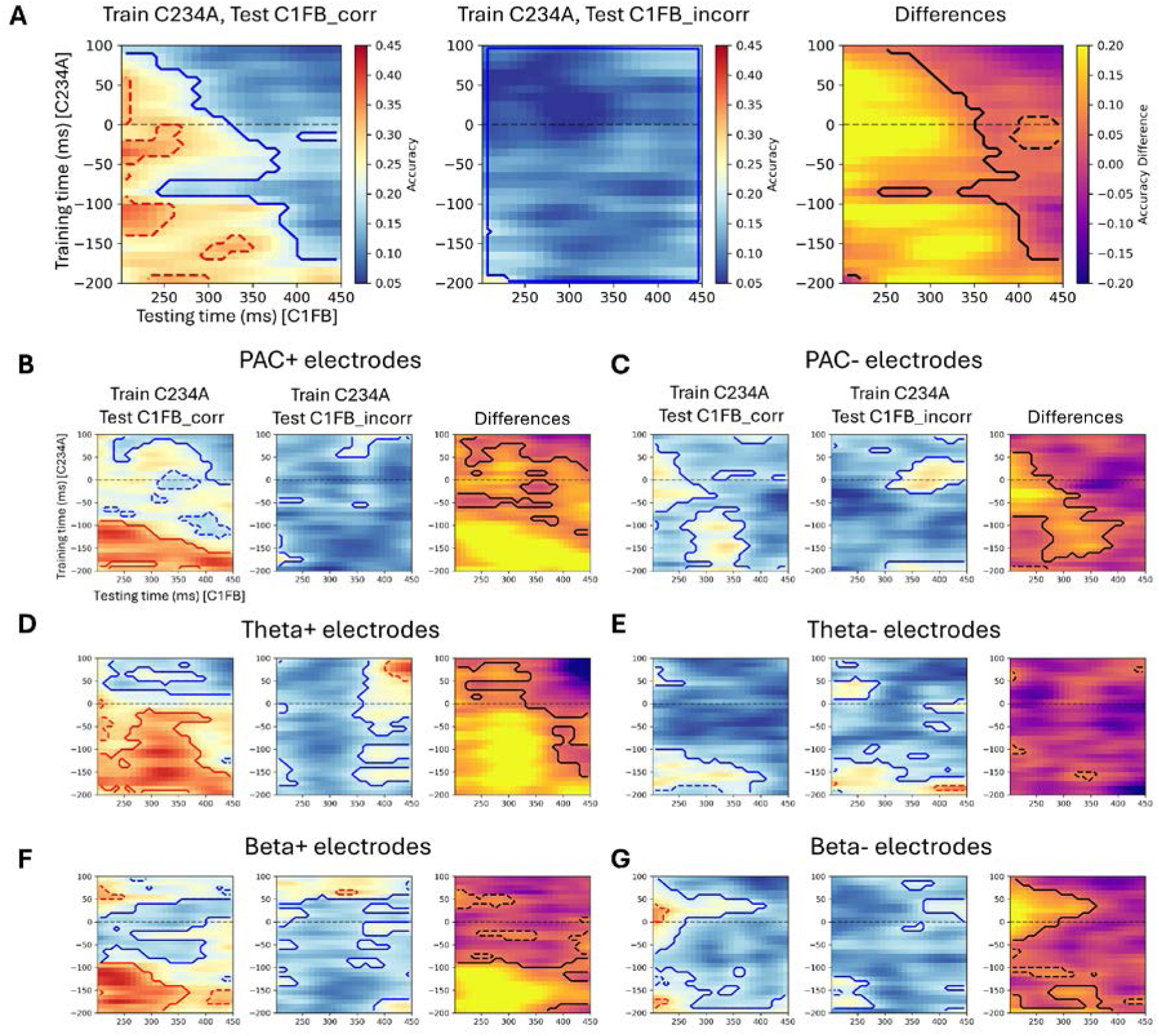
Cross-period generalisation of target object representations and its modulation by oscillatory dynamics. (A) Cross-period generalisation based on local field potentials (LFPs) from all vlPFC electrodes. Classifiers were trained on peri-array neural activity during the retrieval phase (C234A; −200 to 100 ms relative to array onset, y-axis) and tested on the feedback period of the initial learning cycle (C1FB; 200 to 450 ms relative to feedback onset, x-axis). Panels display decoding accuracy for classifiers generalized to correct feedback (C1FB_corr, left) and incorrect feedback (C1FB_incorr, middle) states, as well as their direct difference (right). All cross-period generalisation analyses were performed separately for the two colour-defined object sets and subsequently averaged. Prior to decoding, the centroid amplitude was subtracted at each time point within every trial to isolate object-specific representational structures. (B, C) Cross-period generalisation matrices for vlPFC electrodes showing the strongest (PAC+, B) or weakest (PAC−, C) theta-HFA phase-amplitude coupling (PAC) during 200-450 ms after correct feedback. (D, E) Generalisation matrices for electrodes with the highest (Theta+, D) or lowest (Theta−, E) theta power during 200-450 ms after correct feedback. (F, G) Generalisation matrices for electrodes with the highest (Beta+, F) or lowest (Beta−, G) beta power during 200-450 ms after incorrect feedback. The classification and sorting criteria for all electrode groups (PAC+/−, Theta+/−, and Beta+/−) are identical to those used in Figure 5. For all panels, solid contours denote regions with statistically significant cross-generalisation against chance level (0.25) or significant condition differences (corrected *p* < 0.05, cluster-based permutation test). Dashed contours indicate regions with uncorrected significance (*p* < 0.05). Tests against chance were two-tailed, whereas tests for condition differences were one-tailed.

We next investigated whether this feedback-dependent cross-generalisation was gated by the local oscillatory environment. Again, we grouped the vlPFC electrodes into high (+) and low (−) groups for each oscillatory feature (i.e., theta, beta, and PAC), using the same classification criteria as described for Figure 5. While the cross-generalisation using all electrodes (Figure 6A) yielded relatively modest above-chance clusters for the correct feedback state, focusing on these functionally engaged neural populations revealed a substantially stronger and more widespread alignment. Strikingly, the successful cross-period generalisation and the strong difference between positive and negative feedback were predominantly driven by the dynamically engaged neural populations. Specifically, classifiers within PAC+ (Figure 6B), Theta+ (Figure 6D), and Beta+ (Figure 6F) electrode groups successfully cross-generalized to the correct feedback state and concurrently exhibited widespread, significant condition differences. Notably, within these active functional groups, the robust above-chance decoding for C1FB_corr was prominently concentrated during the pure retrieval preparation phase prior to array onset (training time < 0 ms). In contrast, PAC− (Figure 6C) and Beta-(Figure 6G) electrode groups showed substantially weaker or largely non-significant generalisation differences, and the Theta− group (Figure 6E) exhibited a markedly attenuated effect compared to its Theta+ counterpart. These results provide compelling evidence that the reorganisation of object representations into retrieval-compatible states is not a uniform global process, but rather a targeted transformation actively facilitated by specific rhythmic coordination, namely, elevated PAC and theta activity following positive feedback, and beta activity following negative feedback.

## Discussion

Learning from feedback requires transforming brief outcome signals into stable neural formats that can be reinstated during subsequent memory-guided behaviour. Our results suggest that the vlPFC implements this transformation by reorganizing population geometry in a feedback-dependent manner (Figure 7). Feedback onset elicited robust spiking and LFP responses, with HFA tracking the population firing peak and ERPs differentiating outcome valence. After correct feedback, aperiodic-corrected time-frequency analyses revealed increased theta and a selective rise in theta-HFA PAC; after incorrect feedback, beta power was stronger. Notably, while these macroscopic feedback signatures were robust, they emerged from a functionally heterogeneous population of single neurons and electrodes exhibiting mixed response directions. Despite comparable object information being represented under both kinds of feedback in the vlPFC, multivariate decoders trained within one feedback state consistently outperformed between-feedback generalisation, indicating distinct, state-specific coding formats. Critically, in the vlPFC, population representations sampled after correct feedback were geometrically closer to those reinstated during later retrieval, whereas representations after incorrect feedback diverged from the retrieval geometry. Furthermore, cross-period generalisation analysis directly confirmed this alignment, demonstrating that retrieval-phase neural codes successfully generalized to correct feedback states but dropped below chance during incorrect feedback. Finally, sites with stronger theta power and PAC (post-positive) or stronger beta (post-negative) showed larger modulation of these representational distances and/or cross-period generalisation, suggesting that oscillatory coordination regulates how feedback reshapes object codes toward (or away from) retrieval-relevant states. By contrast, while the dlPFC and TE also exhibited widespread univariate feedback-related responses, they lacked the robust absolute strength of theta-HFA PAC during positive feedback, as well as the stable post-feedback object decoding and state-dependent representational formatting uniquely observed in the vlPFC.

**Figure 7.**
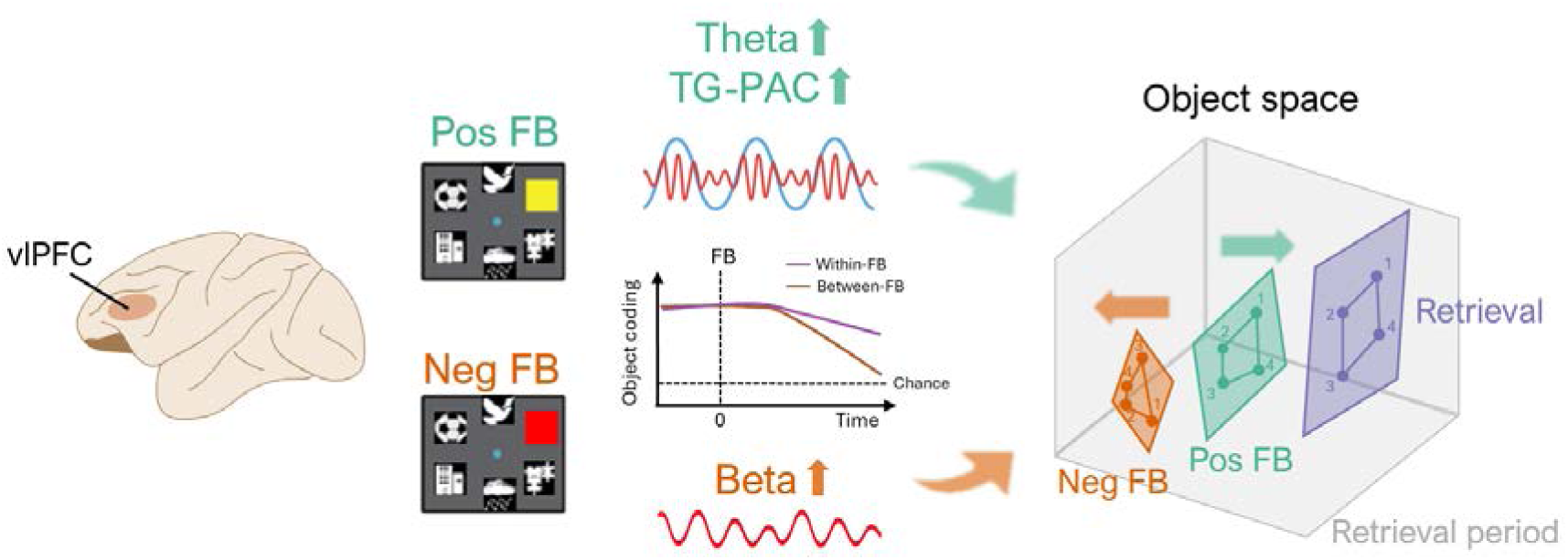
Possible mechanisms underlying learning from feedback in vlPFC. In vlPFC, positive (Pos FB) and negative (Neg FB) feedback elicited distinct neural activity including enhanced theta power and theta-HFA PAC after positive feedback and elevated beta power after negative feedback. Although object information was represented under both feedback conditions, classifiers trained and tested within the same feedback state (within-FB) outperformed those trained across states (between-FB), indicating state-dependent representational formats. In population object space, representations following positive feedback were geometrically closer to those reinstated during later retrieval, whereas negative-feedback states diverged from the retrieval geometry. Also, cross-period generalisation further supported this functional alignment: classifiers trained on the retrieval phase successfully generalized to positive feedback states but dropped below chance during incorrect feedback, indicating an active repulsion from retrieval-compatible format. This feedback-dependent convergence and divergence of representational geometry and cross-period generalisability is further modulated by post-feedback theta-HFA PAC and beta activity. Together, these results suggest that feedback reorganizes object representations in vlPFC through oscillatory coordination, aligning correct-feedback states with future memory retrieval.

Our findings build upon and significantly extend previous descriptive work (*22*), which established the vlPFC as a domain-general hub that represents various kinds of task-relevant information across retrieval phases (Cycles 2-4). While that study focused on identifying the functional role of different brain regions during retrieval, the present work addresses a distinct scientific question: how feedback signals during the initial learning phase (Cycle 1) actively reorganize these representations to support future behaviour. By shifting the focus to the learning phase and incorporating LFP-based oscillatory and cross-frequency coordination, we demonstrate that these rhythmic mechanisms, rather than simple spikes alone, regulate the feedback-driven transformation of neural codes. This revealed dynamic infrastructure provides a physiological bridge between immediate outcome evaluation and the formation of stable, retrieval-compatible representations in the vlPFC.

These findings situate feedback as a driver of dynamic recoding in vlPFC: different feedback signals reconfigure the coding format rather than merely scaling a static code. This view resonates with dynamic coding models that prefrontal neurons continuously adjust their selectivity as behavioural demands evolve (*29–33*), but adds a geometric and functional link to memory: positive feedback not only updates neural activity in the moment but also reorganizes the representational landscape in preparation for future retrieval. In geometric terms, positive feedback appears to reshape population trajectories so that the manifold occupied during learning approaches the subspace later engaged during memory-guided selection. Crucially, our cross-period generalisation results provide an even more powerful and direct demonstration of this format alignment than state-space geometric distances alone. By showing that a classifier trained on the retrieval phase can decode object identity during correct learning feedback (even after removing the overall centroid shift), we demonstrate that positive feedback actively reformats local target codes into a functionally compatible structure that can be directly read out by the downstream retrieval architecture. Conversely, the below-chance generalisation observed during negative feedback implies an active repulsion mechanism, where incorrect outcomes format the population representation into an opposing representational format, effectively shielding the system from consolidating erroneous target associations. Such alignment and anti-alignment could facilitate the reinstatement of relevant target codes while minimizing interference from previously active states. This view parallels recent proposals that learning involves a gradual reorganisation of representational geometry that supports successful behaviour (*31, 57–59*). In this framework, feedback may serve as a teaching signal that projects future retrieval demands onto the current neural space, thereby linking outcome evaluation to the formation of stable and task-relevant representations.

The emergence of this alignment was closely tied to distinct oscillatory signatures, suggesting that rhythmic coordination provides a mechanism for stabilizing the reconfigured representational geometry (*34, 35*). Positive feedback selectively enhanced theta power and theta-HFA PAC, whereas negative feedback increased beta activity. These frequency-specific patterns may suggest complementary computations through which feedback regulates population dynamics. Theta oscillations have been widely linked to adaptive control and updating of internal models (*27, 39–42*), while beta rhythms are thought to stabilize the currently active representational state and suppress competing ensembles (*24, 38, 43*). The concurrent increase in PAC after positive outcomes implies a hierarchical coordination in which slow control rhythms gate fast local assemblies that encode object information (*44–49*). Such cross-frequency coupling may serve as a temporal bridge that binds transient outcome signals to longer-lasting representational formats, supporting credit assignment across successive task epochs. In this framework, oscillations do not merely mark feedback processing but implement a dynamic infrastructure that reconciles flexibility and stability, allowing prefrontal circuits to reorganize representations while preserving continuity in the learned code (*24, 58*).

The unique functional profile of the vlPFC in our task, specifically its ability to implement feedback-driven geometric reorganization through oscillatory coordination, underscores its role as a critical integrative hub supporting learning processes. While widespread neural responses to feedback were observed across the dlPFC and TE, the vlPFC was the exclusive site where these signals were coupled with stable target representation and state-dependent reformatting. This regional specificity aligns with previous work demonstrating that the vlPFC acts as a domain-general hub, encoding various task features more strongly and earlier than dlPFC and TE (*22*). In contrast to area TE, which primarily serves as a sensory provider for object identity, or the dlPFC, which often prioritizes spatial control and executive monitoring (*19, 20, 60, 61*), the vlPFC may serve as an “adaptive coding” engine that bridges learning and retrieval cycles (*22, 33*). Ultimately, the vlPFC uniquely couples outcome evaluation with stable object maintenance, enabling it to actively reformat the representational landscape for future behavioural flexibility (*57*).

Together, our findings reveal how feedback simultaneously shapes local neural dynamics and large-scale representational geometry in the primate prefrontal cortex. In vlPFC, feedback converts transient outcome signals into stable, retrieval-compatible population codes through coordinated oscillatory mechanisms. Positive feedback promotes convergence of object representations toward future retrieval states, supported by enhanced theta-HFA coupling, whereas negative feedback induces divergence accompanied by elevated beta activity. These rhythmic processes provide a dynamic infrastructure that balances flexibility and stability, allowing neural populations to update representations while preserving continuity in the learned code. Conceptually, this study outlines a unified framework in which feedback, oscillatory coordination, and representational geometry jointly implement a mechanism for adaptive cognition. Such principles may generalize beyond feedback learning, describing how prefrontal circuits continually restructure information to sustain flexible, goal-directed behaviour.

## Methods

### Subjects

Two adult male rhesus monkeys (*Macaca mulatta*) participated in the experiments (∼14 kg each). All experimental procedures complied with the Animals (Scientific Procedures) Act 1986 of the UK and were approved by the Home Office under a Project License following review by the University of Oxford Animal Care and Ethical Review Committee. Procedures conformed to the European Community guidelines for the care and use of laboratory animals (EUVD, European Union directive 86/609/EEC).

### Experimental Task

Monkeys performed a multi-cycle object-learning task in which they learned and selected visual targets for soft food rewards. The task required animals to discover target objects through trial-and-error feedback in the first learning cycle and subsequently retrieve the same targets based on colour cues in later cycles (Figure 1A). Each recording session contained a series of problems; each composed of four cycles of trials. On average, monkeys completed approximately 57 problems per session, all using a fixed set of eight objects. Four of these objects were associated with a green cue (Figure 1A, inset), and four with a yellow cue. Six objects were used in each problem, excluding the two target objects from the preceding problem.

Before each trial began, a central white fixation point (FP) appeared together with six surrounding black squares (6 × 6° visual angle, centered 14° from fixation). To initiate the trial, the animal was required to fixate on the FP within a window of 2.6 × 2.6° (animal A) or 3 × 3° (animal B). Upon fixation, the FP turned red, initiating a short preparatory delay (0.3-0.6 s). A cue stimulus (6 × 6° square) then appeared at the fixation point. In the initial learning cycle (Cycle 1), the cue was grey, providing no information about which target to select; in subsequent retrieval cycles (Cycles 2-4), the cue was either green or yellow, indicating which of the two previously rewarded targets should be chosen.

After a fixed delay of 0.5 or 1 s (fixed for each session), the circular array of black placeholders was replaced by six choice objects. Following a variable delay of 1 to 2.5 s, the FP changed to cyan, serving as the “go” signal. The cue stimulus simultaneously disappeared, and the animal was required to make a saccade to one of the objects within 1 s. After the chosen object was fixated and held for 0.35-0.45 s, it was replaced by a visual feedback signal (FB), presented as a 6 × 6° square.

Positive feedback consisted of the appropriate cue colour (green or yellow) and remained visible for 0.4 s. This was followed by a brief delay of 0.05–0.15 s (resulting in a total delay of 0.45-0.55 s from feedback onset to reward delivery), after which a drop of soft food reward was delivered and the inter-trial interval (ITI) commenced. Negative feedback was indicated by a red square and was not rewarded. For standard trials within a cycle, the ITI display (consisting of a central white FP with surrounding black squares) was maintained for 0.7–0.9 s. Consummatory licking was neither constrained nor explicitly measured, and actual reward consumption primarily occurred during the ITI period.

Distinct displays indicated transitions between task phases. To signal the end of a cycle, the standard ITI was preceded by a brief period showing only a grey FP (3.2-3.5 s), while at the end of a problem, the screen blanked for 3.3-3.6 s before a new pair of targets was introduced. To initiate any new trial, including those following an aborted or non-rewarded trial, the monkey was required to establish fixation on the white FP, which appeared as a discrete trigger to signal trial availability. Trials were aborted without reward and excluded from analysis if fixation was broken before the go cue or if gaze was not maintained until feedback onset.

All task events, including visual presentation, timing control, and reward delivery, were managed using the REX real-time data acquisition and control software (Laboratory of Sensorimotor Research, NIH). Stimuli were presented on a 17.5-inch LED screen positioned in front of the animal. Eye position was continuously monitored at 120 Hz using an infrared eye-tracking system (Applied Science Laboratories) and synchronized with neural and behavioural event markers.

### Neural Recordings

Each monkey was implanted with a titanium head holder and recording chambers (form-fitting chamber system, Gray Matter Research) under general anesthesia using aseptic surgical procedures. Frontal chambers were positioned over the right lateral prefrontal cortex in both animals (monkey A: anterior-posterior (AP) = 34.9 mm, medio-lateral (ML) = 14.7 mm; monkey B: AP = 36.4 mm, ML = 19.1 mm). A second chamber was placed over the right TE (monkey A: AP = 4.3 mm, ML = 11.3 mm; monkey B: AP = 3.0 mm, ML = 18.3 mm). Craniotomies were made under each chamber to permit electrophysiological recordings.

Neural signals were collected using semi-chronic microdrive arrays (SC-96 for frontal cortex, SC-32 for TE; Gray Matter Research) with 1.5 mm inter-electrode spacing. Electrodes were independently movable between sessions, enabling recordings from largely non-overlapping neural populations across days. The frontal array targeted extensive regions of the lateral prefrontal cortex encompassing vlPFC and dlPFC areas on the cortical convexity, while the temporal array covered ventral subdivisions of TE.

Neural activity was recorded at 30 kHz, amplified, and band-pass filtered (300 Hz to 10 kHz) using a multichannel recording system (Cerebus, Blackrock Microsystems) and stored for offline spike sorting (Offline Sorter, Plexon).

Local field potentials (LFPs) were simultaneously acquired from the same electrodes, low-pass filtered at 250 Hz, and down-sampled to 1 kHz for analysis. To remove electrical line noise, the continuous LFP signals were notch-filtered at 50 Hz and its harmonics (100, 150, and 200 Hz). The data were then epoched around the events of interest. To eliminate motion and electrical artefacts, we applied a strict thresholding procedure: for each electrode, the voltage traces were z-scored across time, and any trial containing fluctuations exceeding 6 standard deviations from the mean was excluded. Furthermore, to ensure robust signal quality, if an electrode exhibited such artefactual responses in >20% of the trials within a given session, it was excluded from all subsequent LFP analyses.

Between sessions, electrodes were advanced by at least 62.5 µm to ensure sampling of new units. Recordings were conducted over a total of 247 daily sessions (122 with monkey A and 125 with monkey B). No pre-selection of neurons based on task responsiveness was performed; recordings proceeded once well-isolated single-unit activity was obtained.

At the completion of experiments, animals were deeply anesthetized with barbiturate and perfused transcardially with heparinized saline followed by 10% formalin. Brains were removed for histological verification of recording sites, which confirmed electrode tracks within the intended regions.

### Univariate Analyses on Spiking Activity and LFPs

To characterize feedback-related neural responses in vlPFC, dlPFC, and TE, we analysed both spiking activity and LFPs aligned to feedback onset during the learning cycle (Cycle 1).

#### Population firing rate

Spiking activity was smoothed with a Gaussian kernel of 50 ms full-width-at-half-maximum (FWHM) and converted to firing-rate time series aligned to feedback onset. For each neuron, firing rates were z-scored across all trials and time points and averaged separately for correct and incorrect feedback. Population activity was then obtained by averaging normalized firing rates across neurons, producing mean time courses that captured the temporal profile of feedback-locked population responses. To control for unequal trial counts, we also re-ran statistical tests on a subsampled dataset where correct and incorrect trials were randomly matched to a strict 1:1 ratio for each object condition within every session.

#### High-frequency activity (HFA)

To estimate high-frequency activity associated with local spiking, LFPs were filtered in the 100-250 Hz range and transformed using the analytic amplitude of the Hilbert signal. Instantaneous power (squared amplitude) was then smoothed with a Gaussian kernel (FWHM = 50 ms) and then downsampled to 100 Hz for analysis. All signals were aligned to feedback onset and z-scored within each electrode across all trials and time points. As in other univariate analyses, for each electrode, HFA time courses were averaged across trials within each feedback condition (correct vs. incorrect) and then averaged across electrodes.

#### LFP voltage analyses

Time-domain LFP activity (i.e., ERP) was low-pass filtered below 20 Hz to extract slow potential fluctuations, down-sampled to 100 Hz, and aligned to feedback onset. Data were z-scored within each session and averaged across electrodes to obtain regional ERPs for correct and incorrect feedback. The same 1:1 trial-matching subsampling control was applied to LFP voltage signals to verify the robustness of these feedback-related dynamics.

#### FOOOF and oscillatory power analyses

To obtain aperiodic-adjusted oscillatory power time series, we combined time-frequency decomposition with spectral parameterization using the FOOOF algorithm (*51*). Broadband LFPs were first band-filtered to 2-30 Hz and transformed using Morlet wavelets with frequency-dependent cycles (*n_cycles* = frequency/2). Prior to decomposition, phase-locked ERP activity for each object condition was subtracted from each trial to remove non-oscillatory slow components. For each electrode, trial, and time point, the resulting power spectra were parameterized using FOOOF in a fixed aperiodic mode (frequency range 2-30 Hz, peak width limits [2, 8] Hz, peak threshold 2.0, and maximum of 4 peaks). The aperiodic component (1/f background) was subtracted from the total spectrum to yield periodic, oscillatory power estimates. This procedure produced time-resolved, aperiodic-adjusted oscillatory power matrices for each recording site. Power values were z-scored within each electrode across trials and time points and down-sampled to 100 Hz for analysis. Oscillatory power in the theta (4-8 Hz) and beta (20-30 Hz) bands was then extracted from the FOOOF-corrected spectra and compared between correct and incorrect feedback conditions.

#### PAC analyses

To quantify theta-HFA phase-amplitude coupling (PAC) following feedback, we filtered LFPs into two frequency ranges and then computed the modulation index (MI) (*53*). Specifically, we defined a low-frequency range of 2-30 Hz sampled every 2 Hz and a high-frequency range of 40-250 Hz sampled every 8 Hz. For each low/high pair, we band-pass filtered the signal, extracted phase (low band) and amplitude envelope (high band) with the Hilbert transform, and estimated the MI using the Tort method (*53*), as implemented in the toolbox *pactools* (*62*). We aligned data to feedback onset, subtracted the condition-specific ERP to suppress slow potentials, and restricted PAC estimation to the 200-450 ms post-feedback window. To normalize MI and control for spurious coupling, we built a surrogate distribution by circularly time-shifting each single-trial time series (200 iterations), which preserves spectral content while disrupting cross-frequency dependencies. We then converted observed MI to z-scores at every frequency pair using:

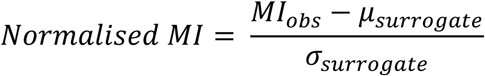

These surrogate-corrected normalised MI maps were averaged across electrodes and compared between correct and incorrect feedback conditions.

To further validate the frequency specificity and robustness of the coupling, we conducted two control analyses using the same surrogate-correction procedure. First, to dissect the specific high-frequency components modulated by theta phase, we split the HFA range and computed theta-HFA PAC separately for a lower amplitude band (100–150 Hz) and an upper amplitude band (175–250 Hz). Second, to test whether the feedback-induced theta and beta rhythms reflect nested or independent processes, we computed theta-beta PAC specifically using theta phase (4–8 Hz) and beta amplitude (20–30 Hz).

### Functional heterogeneity of feedback responses across neurons and electrodes

To evaluate the functional heterogeneity of feedback-related modulations across the population of vlPFC, dlPFC, and TE, we examined single-trial responses for each individual neuron and electrode. Specifically, for firing rate, LFP voltage, HFA, and oscillatory theta/beta power, we computed the mean activity within the 200–450 ms feedback window on a trial-by-trial basis. We then compared the distribution of single-trial responses between correct and incorrect feedback conditions using an independent samples Welch’s t-test (assuming unequal variances). For theta-HFA PAC, however, because single-trial phase-amplitude estimation inherently suffers from a low signal-to-noise ratio, we instead employed a non-parametric trial-label permutation approach to maintain high signal-to-noise ratio while assessing statistical significance. For each electrode, we randomly shuffled the condition labels 200 times (balancing computational efficiency and statistical validity) to compute normalised PAC differences (with 200 surrogates), thereby constructing an empirical null distribution. Based on an alpha level of 0.05, each neuron or electrode was classified into one of three categories: (1) significant positive modulation (Correct > Incorrect), (2) significant negative modulation (Correct < Incorrect), and (3) non-significant. This procedure allowed us to quantify the exact proportion of neurons and electrodes exhibiting response polarities that matched, opposed, or were insensitive to the population-level average.

### Multivariate Decoding Analyses

To decode object-specific neural representations during the feedback period (Figure 3), we used data from all recorded neurons (for spike-based decoding) or electrodes (for LFP-based decoding) across sessions and applied time-resolved multivariate pattern analysis. Data were aligned to feedback onset, down-sampled to 100 Hz, and normalized within each neuron/electrode by z-scoring across all trials and time points. We implemented a bootstrapped pseudotrial approach to improve signal-to-noise while maintaining trial-level variability. For each object condition, all trials were randomly split into two independent halves for training and testing. Within each half, we performed 30 bootstrap resampling iterations; in each iteration, trials were resampled with replacement and averaged to generate one pseudotrial. Consequently, each half contained 30 pseudotrials per object condition, yielding 120 pseudotrials in total for each object set (4 objects × 30 pseudotrials). The training set therefore consisted of 120 pseudotrials from one half, and the test set of 120 pseudotrials from the other.

Decoding was conducted separately for the two object sets (green and yellow), each comprising four objects (chance level = 0.25). A linear support vector machine (SVM) classifier was trained to discriminate among the four objects within each set. Input features consisted of concatenated firing rates (or LFP amplitudes) across all neurons/electrodes at each time point. To increase the signal-to-noise ratio for each decoding sample, neighbouring timepoints (±2 samples) were concatenated as additional features. Prior to classification, features were whitened and reduced using PCA (variance retained = 95%). The classifier was trained and tested on independent pseudotrial sets (training set vs. testing set) for within-feedback decoding (e.g., correct → correct, incorrect → incorrect) and between-feedback generalisation (e.g., correct → incorrect, incorrect → correct). Decoding accuracy was computed at each time point, producing time-resolved accuracy curves that were averaged across repetitions. The entire procedure was repeated 100 times to obtain the final population-level decoding curves.

### Representational Geometry Analyses

To examine the structure of object representations across learning and retrieval, we constructed a population activity space using PCA based on neural population activity from the peri-array period of Cycles 2-4 (−200 to 100 ms from array onset; C234A). We then examined projections into this space from three task periods: Cycle 1 feedback after correct trials (200-450 ms from feedback onset; C1FB_corr), Cycle 1 feedback after incorrect trials (200-450 ms from feedback onset; C1FB_incorr), and C234A itself. Analyses were performed separately for the two object sets and for both spiking activity and LFP voltage. The Cycle 1 feedback window was restricted to 200–450 ms post-feedback.

The 200 ms onset was chosen to bypass early sensory-evoked, phase-locked responses and to allow sufficient time for stable outcome-evaluation signals to reach the vlPFC. The 450 ms endpoint was constrained by the task design to ensure no contamination from stimulus offset, as feedback duration was jittered between 450 and 550 ms. For the retrieval state (Cycles 2-4), we selected a peri-array window of −200 to 100 ms relative to array onset. Based on previous analyses showing that target encoding in the vlPFC emerges during the delay period from ∼ 300 ms after the cue onset (*22*), and given that our cue-to-array delay was either 0.5 or 1 s depending on the session, a window starting 200 ms prior to array onset consistently captures the late delay period when these target signals have already stabilized. This window was designed to capture the internally driven reinstatement of the target object in memory and its transition into the early phase of array processing to guide memory-guided selection.

For each neuron or electrode, activity values were averaged within the relevant time windows, down-sampled to 100 Hz, and normalized by z-scoring across all trials and time points within a session. Correct and incorrect trials were further balanced by serial position (see Statistical analyses for details) and object identity to ensure matched sampling across conditions.

To construct the population vectors, neural responses for each of the eight objects were averaged across trials and randomly split into two independent halves (Split A/B) to allow cross-validation. For each neuron, Split A and B means formed two independent representations per object. We concatenated these across all neurons (or electrodes) to form population activity matrices of dimension *neurons × objects × splits*. We then applied PCA separately for each object set (four objects per set) to reduce dimensionality to the first three components.

We trained the PCA model on neural population activity from the retrieval period (C234A) using one independent data split (Split A), and projected activity from all periods (C1FB correct, C1FB incorrect, and C234A) from the held-out split (Split B) into this retrieval-based space. This cross-validated approach ensured that the PCA projection and subsequent geometric analyses were performed on independent data. The resulting 3-D object constellations for each condition were visualized and compared within this common representational space.

To quantify representational relationships, we generated bootstrapped samples (1000 iterations) by resampling neurons (or electrodes) with replacement within one split (Split B) to obtain variability estimates. Each bootstrap sample was projected into a fixed PCA space (trained on C234A), and distances were computed between the feedback-related object constellations (C1FB correct or incorrect) and the retrieval constellation (C234A). Centroid distances were computed as the Euclidean distance between the mean positions of the four-object constellations across periods. The resulting bootstrap distributions were used to compute 95% confidence intervals and to statistically compare the correct- and incorrect-feedback conditions using paired *t*-tests.

To test how oscillatory activity modulated these geometric effects, we repeated the same analysis using subsets of electrodes grouped by their theta, beta, or PAC strength. Electrode selection was based on the previously described FOOOF-based time-frequency and PAC analyses. For theta power and PAC, we averaged activity during the 200-450 ms post-feedback window after correct feedback, ranked all electrodes by their average strength, and defined the upper and lower 1/3 electrodes as Theta+/Theta− and PAC+/PAC− groups, respectively. For the Theta+/Theta− grouping, because animal A showed a secondary cluster of electrodes with unusually high theta power during correct feedback (see Figure S9), we excluded electrodes exceeding 0.45 power units (corresponding to the local minimum between the two peaks of the distribution) prior to defining Theta+/Theta− groups to ensure consistent ranking across animals. For beta power, electrodes were ranked according to post-feedback activity after incorrect trials, and the upper and lower 1/3 electrodes were taken as Beta+/Beta− groups. We then performed geometry analyses independently within each of these electrode subsets to assess how oscillatory dynamics influenced the stability of object-space organization.

### Cross-period generalisation analyses

To test whether the neural codes formed during learning feedback (Cycle 1) functionally align with those utilized during later memory retrieval (Cycles 2-4) (Figure 6), we performed a cross-period generalisation analysis. To enhance decoding sensitivity to complex neural dynamics, we utilized a recently developed decoding framework: Heterogeneous reservoir computing (HeteroRC) (*56*). Briefly, HeteroRC acts as a powerful non-linear feature extractor that projects multivariate neural time series into a high-dimensional recurrent state space governed by heterogeneous intrinsic time constants. This creates a multiscale temporal filter bank that integrates both fast and slower neural dynamics directly from the raw time series, which are subsequently decoded via a linear ridge-regression readout. For this analysis, we used LFP data down-sampled to 100 Hz. Unlike the analyses above, we did not apply a 20 Hz low-pass filter, allowing the reservoir to exploit broader spectral information. Crucially, to isolate object-specific representational structures and prevent decoding from being driven by global, non-specific signal shifts, we removed the centroid shift prior to decoding. Specifically, at each time point, the overall mean amplitude across all four objects within a set was subtracted from each trial.

Similar to our standard multivariate decoding described above, we employed a bootstrapped pseudotrial approach (30 pseudotrials per object) and performed the decoding separately for the two object sets. The HeteroRC reservoir consisted of 1000 units. To capture a physiologically relevant hierarchy of timescales, the intrinsic time constants of these units were sampled from a log-normal distribution (tau_mode = 0.03 s, tau_sigma = 0.8, tau_max = 100 ms). Other reservoir hyperparameters were kept at their default values (input scaling = 0.5, bias scaling = 0.5, spectral radius = 0.95, bidirectional = False).

For cross-period generalisation, the linear readout of the HeteroRC model was trained on the peri-array neural activity during the retrieval phase (C234A; −200 to 100 ms relative to array onset) and subsequently tested on the feedback period of the initial learning phase (C1FB; 200 to 450 ms relative to feedback onset). This generalisation was evaluated separately for correct and incorrect feedback trials. Finally, to assess the modulatory role of oscillatory dynamics, this cross-generalisation procedure was repeated using data restricted to specific electrode subpopulations (PAC+/−, Theta+/−, and Beta+/−) defined by the same criteria used in the representational geometry analyses.

### Statistical Analyses

We used nonparametric, resampling-based statistics throughout. For both univariate and multivariate decoding analyses, we used permutation methods to construct empirical null distributions while explicitly controlling the potential confound from serial-position effects.

In each learning cycle, monkeys viewed three objects per colour set, resulting in a structured sequence of possible outcomes: the first object (serial position 1) had a 33.3% chance of being correct, the second (serial position 2) 50%, and the third (serial position 3) 100%, as each object could only be selected once and revisits were excluded from analysis. Consequently, animals were likely to anticipate negative feedback early and positive feedback late within a cycle. To remove this confound, permutation of feedback labels was performed within each serial-position bin, preserving the empirical distribution of serial positions while disrupting the mapping between feedback type and neural response.

For each comparison (e.g., correct vs. incorrect feedback), we generated 1000 permutations. At each time point (or each time-by-time bin for cross-period generalisation), we compared the observed difference between conditions against the null distribution obtained from the permuted data to compute pointwise *p* values. We used cluster-based permutation tests (*63*) to control for multiple comparisons across time (or time-by-time bin for cross-period generalisation), and clusters with corrected *p* < 0.05 were considered significant. All tests were two-tailed unless otherwise specified. For the within-between decoding comparison and the cross-period generalisation difference (correct vs. incorrect feedback, we used a one-tailed test based on the a priori hypotheses that within-feedback decoding accuracy would exceed cross-feedback decoding, and that cross-generalisation to the correct feedback state would exceed that to the incorrect state. For the representational geometry analyses, significance of distance metrics was determined from bootstrap distributions (1000 iterations). Confidence intervals were computed directly from the bootstrap samples; effects were considered significant if the 95%, 99%, or 99.9% confidence intervals did not include zero (corresponding approximately to *p* < 0.05, *p* < 0.01, and *p* < 0.001, respectively).

## Acknowledgements

This project was supported by UKRI MRC intramural funding MC_UU_00030/7 to D.J.M and J.D, and MC_UU_00030/15 to A.W. R.L. was supported by a Gates Cambridge Scholarship (OPP1144) and a postdoctoral fellowship from the Canadian Institutes for Health Research (200883). For the purpose of open access, the author has applied a Creative Commons Attribution (CC BY) licence to any Author Accepted Manuscript version arising from this submission.

## Author contributions

M. Kadohisa, M. Kusunoki, M.J.B., and J.D. designed the research. R.L. and J.D. conceived the study. M. Kadohisa and M. Kusunoki collected data. R.L., D.J.M, and J.D. analysed data. R.L. wrote the first draft of the paper. All authors contributed to the final version of the paper. J.D. supervised the work.

## Supplementary

**Figure S1.**
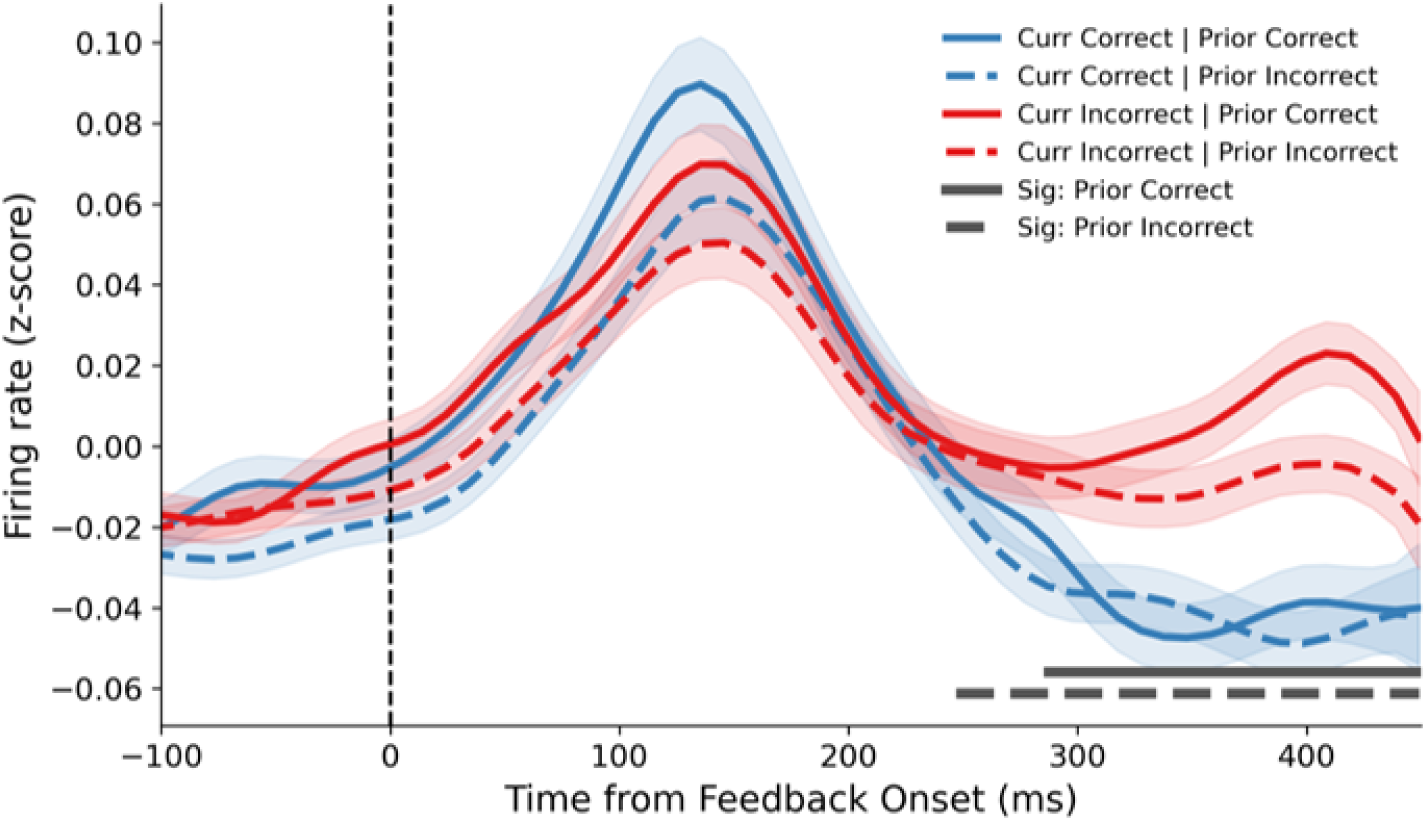
Single-trial-back analysis of population firing rates in vlPFC during the learning cycle. Population firing rates aligned to feedback onset (time 0) are shown for four conditions, defined by the outcome of both the current and the immediately preceding N-1 trial. Trials were categorized as Current Correct given Prior Correct (CC, solid blue), Current Correct given Prior Incorrect (CI, dashed blue), Current Incorrect given Prior Correct (IC, solid red), and Current Incorrect given Prior Incorrect (II, dashed red). Shaded regions denote ±SEM across 1,252 valid neurons. Horizontal solid and dashed grey bars at the bottom indicate time windows with significant differences between current correct and current incorrect feedback under Prior Correct (CC vs. IC) and Prior Incorrect (CI vs. II) conditions, respectively (cluster-corrected *p* < 0.05). Significance was assessed using permutation tests that stringently controlled for both serial positions and prior feedback outcomes. The overlapping significance windows demonstrate that the feedback-driven divergence in vlPFC is robustly determined by the current outcome, independent of prior reward history.

**Figure S2.**
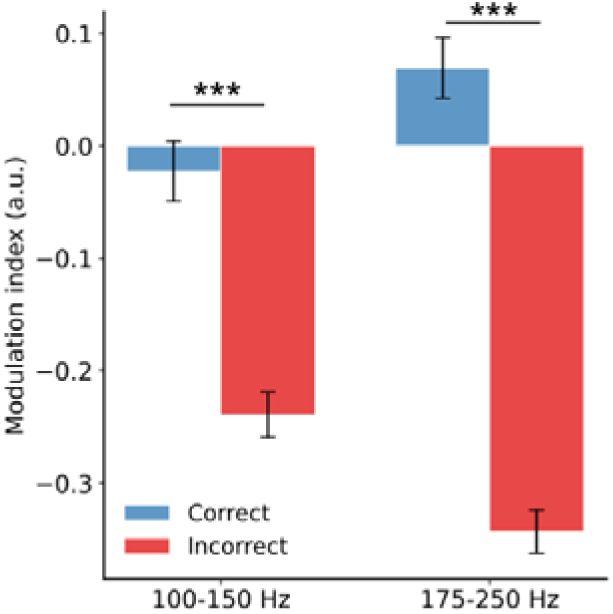
Comparison of PAC results (4-8 Hz phase, 100-150 Hz or 175-250 Hz amplitude) during 200-450 ms window for correct and incorrect feedback (same calculation as Figure 2I, except for different high-frequency range). Error bars denote ± SEM across electrodes. *** *p* < 0.001

**Figure S3.**
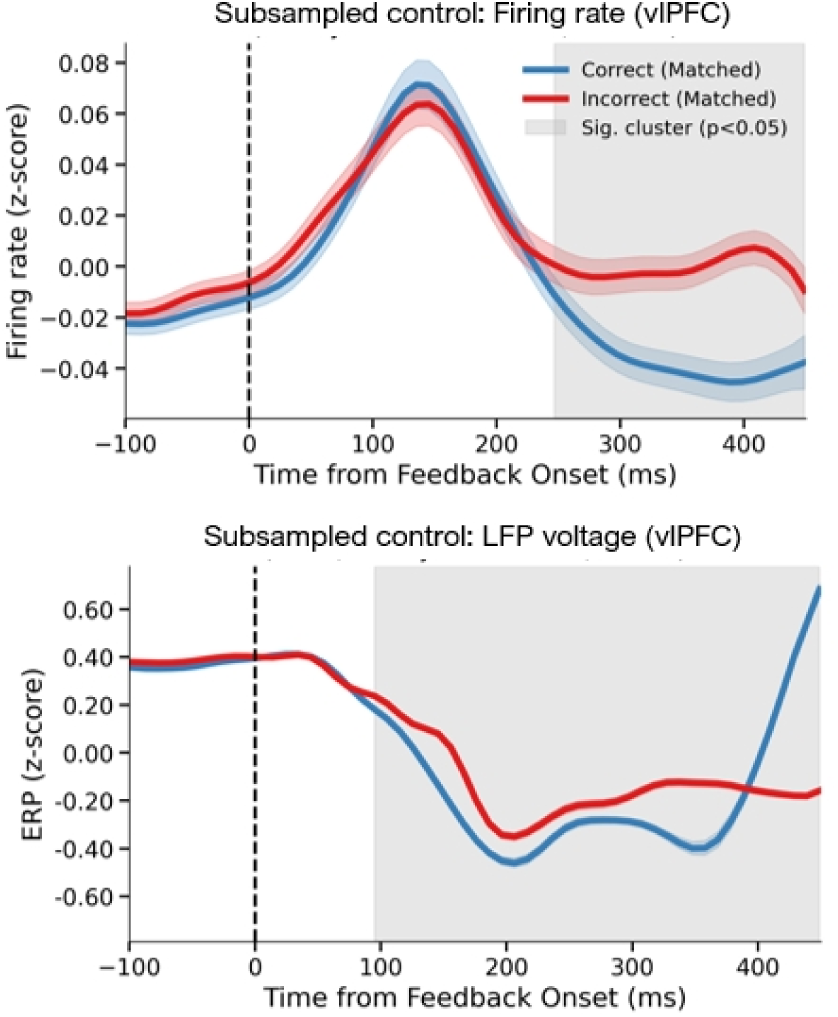
Subsampled control analysis of firing rates and LFP voltages. (Top) Population firing rates and (Bottom) LFP voltages (0-20 Hz) aligned to feedback onset, computed from these balanced subsets for correct (blue) and incorrect (red) feedback. Shaded areas around the curves denote ± SEM. Grey shaded regions indicate time windows with significant differences between conditions (cluster-corrected *p* < 0.05). Significance was established by re-running 1000 permutations explicitly on the subsampled data.

**Figure S4.**
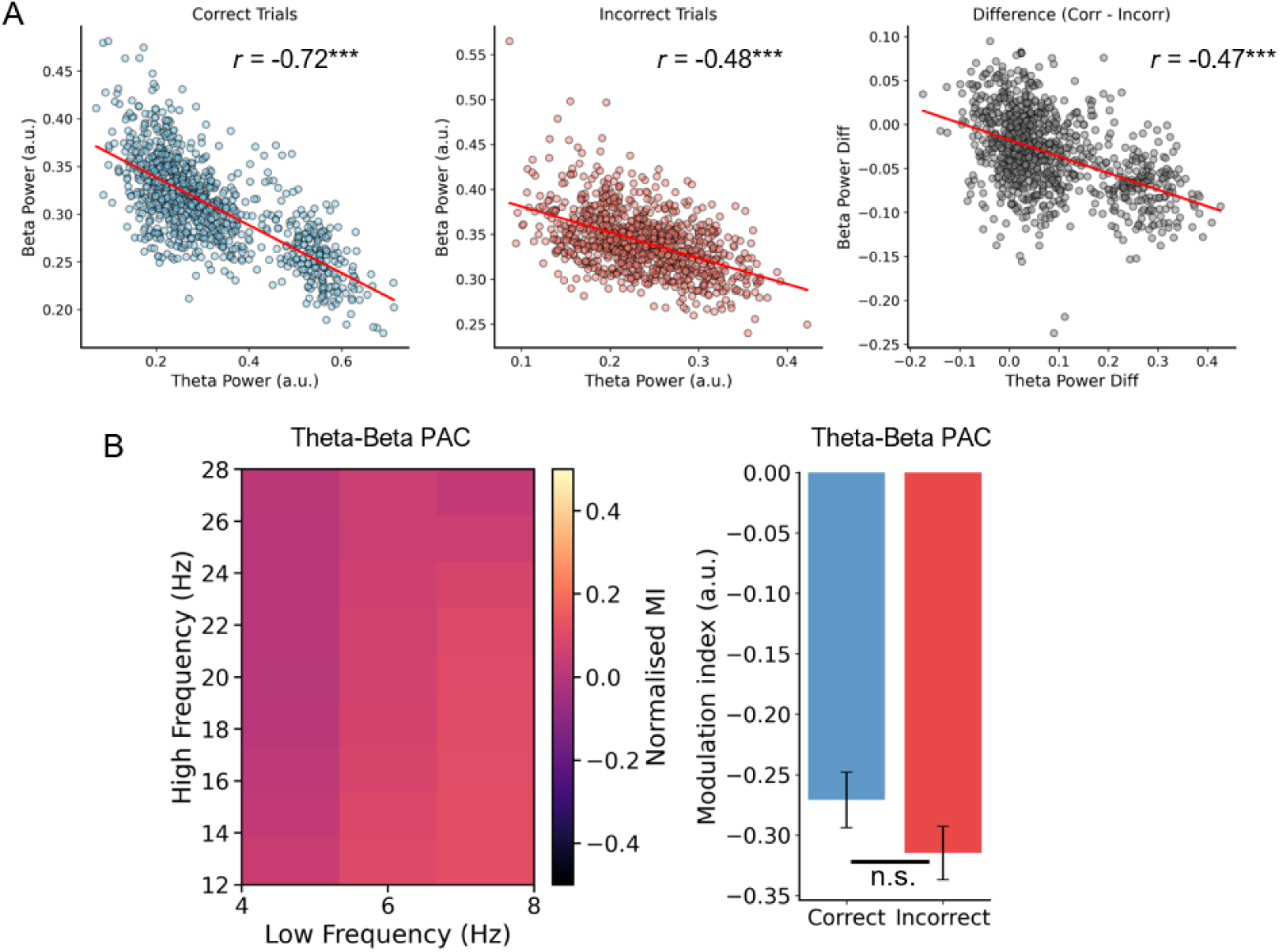
Spatial correlation and cross-frequency coupling between theta and beta oscillations in vlPFC. (A) Spatial correlation between theta (4-8 Hz) and beta (20-30 Hz) power across all recorded vlPFC electrodes during the late feedback window (200–450 ms). Scatter plots show the mean power for correct trials (left), incorrect trials (middle), and the power difference between conditions (right; Correct minus Incorrect). Solid red lines denote linear regression fits. r values represent Pearson correlation coefficients. *** *p* < 0.001. (B) Analysis of theta-beta phase-amplitude coupling during the same 200–450 ms window. Left: Heatmap displaying the difference in normalized modulation index (MI) between correct and incorrect feedback across theta phase and beta amplitude. Right: Comparison of Theta-Beta PAC computed specifically for theta phase (4–8 Hz) and beta amplitude (20–30 Hz) between correct and incorrect trials. Error bars denote ± SEM across electrodes.

**Figure S5.**
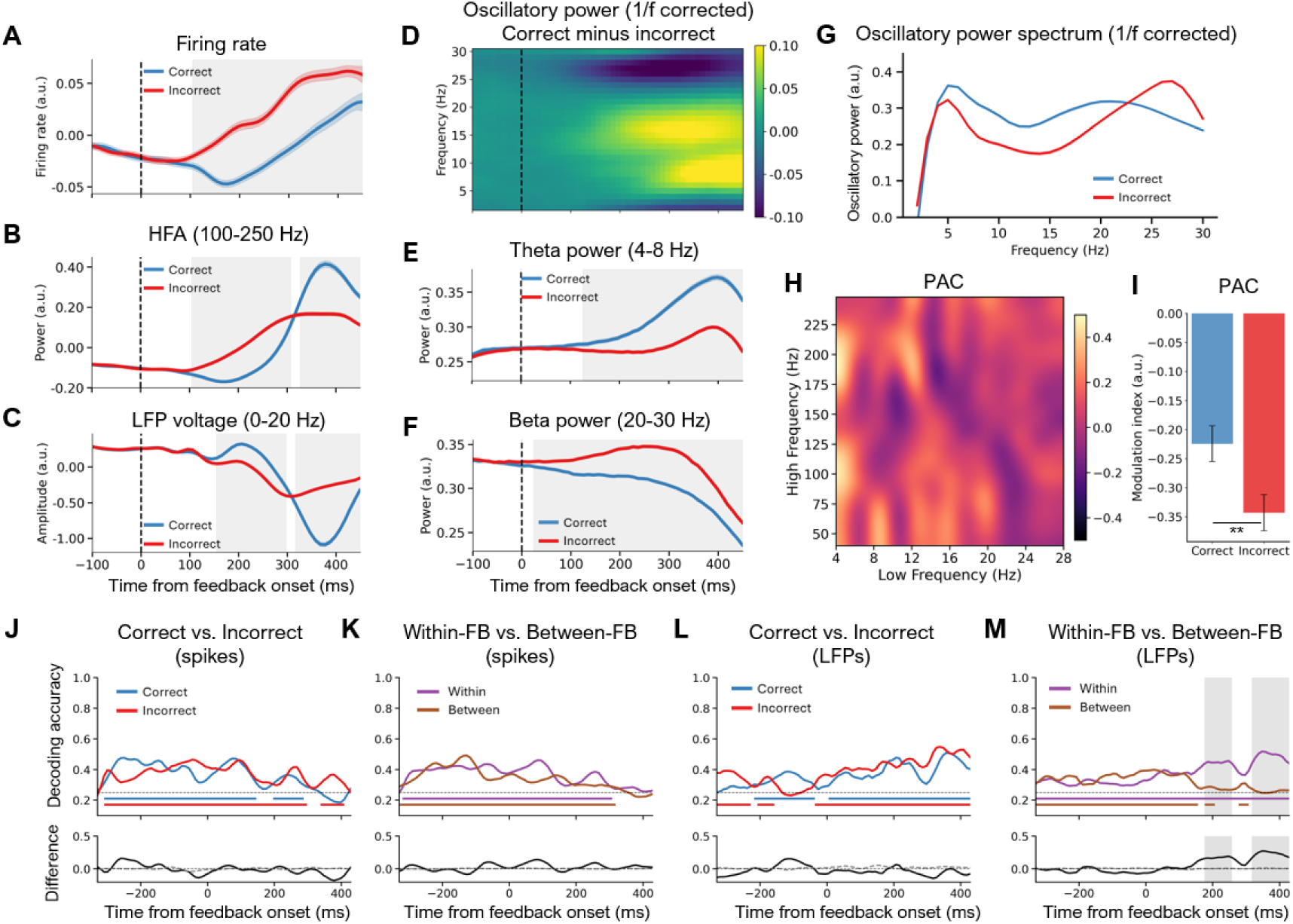
Feedback-related spiking and LFP responses and object representations in dlPFC. (A) Population firing rates averaged across dlPFC neurons for correct and incorrect feedback. (B) High-frequency activity (HFA, 100-250 Hz) averaged across dlPFC electrodes for correct and incorrect feedback. (C) Event-related potentials (ERPs) averaged across dlPFC electrodes for correct and incorrect feedback. (D) Time-frequency decomposition results of oscillatory power (correct minus incorrect feedback conditions) after removing the aperiodic 1/f component using FOOOF. (E-F) Time courses of theta (4-8 Hz) and beta (20-30 Hz) power for correct and incorrect feedback. (G) Aperiodic-corrected power spectra computed using the 200-450 ms post-feedback window. (H) Cross-frequency phase-amplitude coupling (PAC) between low (4-28 Hz) and high (40-250 Hz) frequency ranges, shown as difference between positive and negative feedback. (I) Theta-HFA PAC (4-8 Hz phase, 100-250 Hz amplitude) during 200-450 ms window for correct and incorrect feedback. (J-K) Firing-rate-based decoding of target objects during the feedback period in Cycle 1. (J) Decoding accuracy for correct (blue) and incorrect (red) feedback trials, with the lower panel showing their difference. (K) Decoding accuracy when classifiers were trained and tested within the same feedback condition (within-FB) or across conditions (between-FB), with the lower panel showing within-minus-between differences. (L-M) Same analyses as in (J-K) but based on LFP voltage signals. All analyses were aligned to feedback onset (time 0). Horizontal dotted line indicates theoretical chance level (0.25). Coloured horizontal bars indicate time windows in which decoding accuracy was significantly above chance (corrected p < 0.05, cluster-based permutation test). Shaded regions denote time windows with significant differences between conditions (corrected p < 0.05, cluster-based permutation test). In the difference panels, the grey dashed line shows the permutation-derived baseline (chance-level). Gray shading regions indicate time windows with significant differences between correct and incorrect trials (corrected p < 0.05, cluster-based permutation test). Significance was assessed by constructing empirical null distributions from permutation samples while accounting for serial-position effects. Error shading or error bars denote ± SEM across neurons or electrodes. ** p < 0.01.

**Figure S6.**
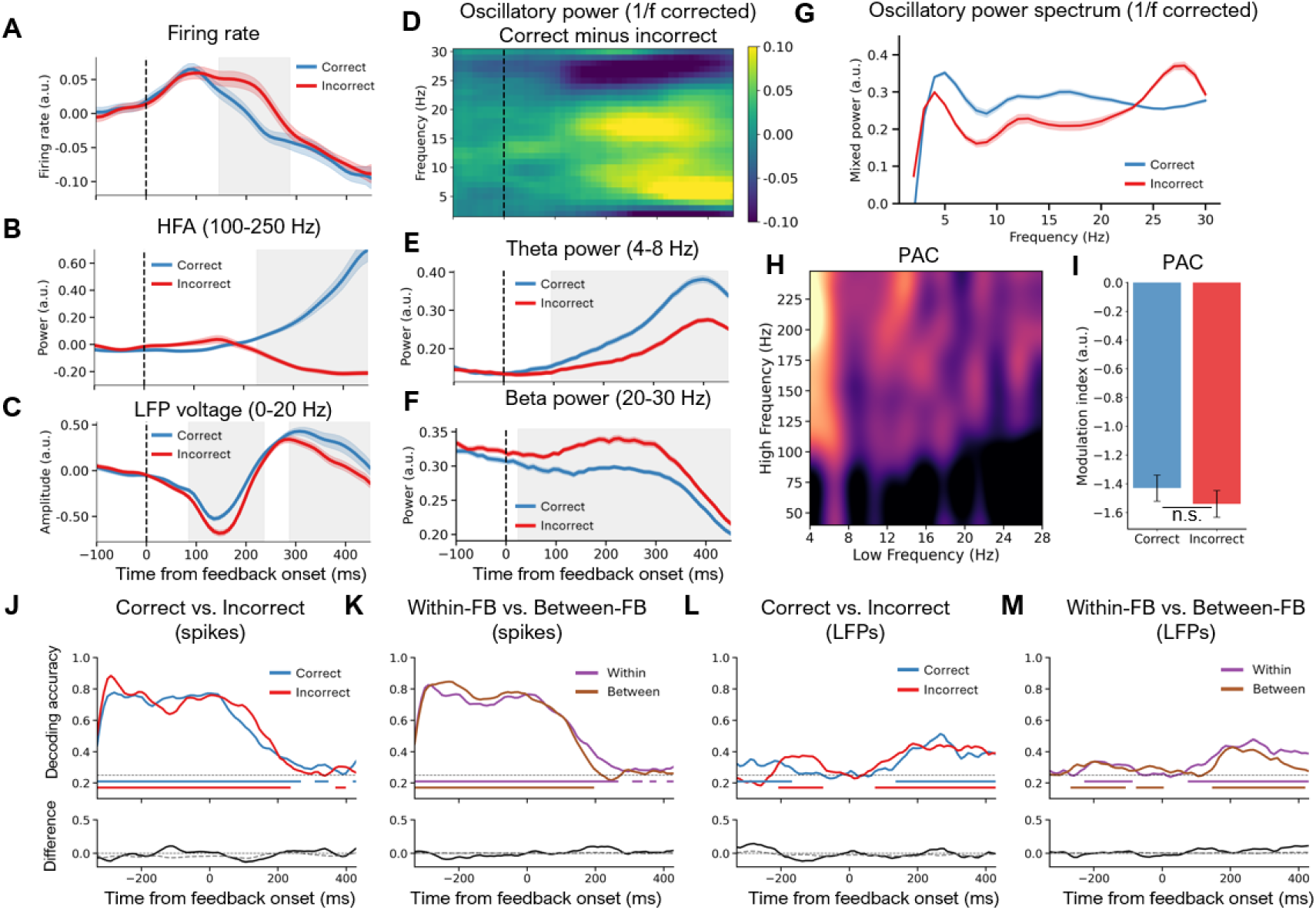
Feedback-related spiking and LFP responses and object representations in TE. Same analyses as in Figure S5 but based on TE signals.

**Figure S7.**
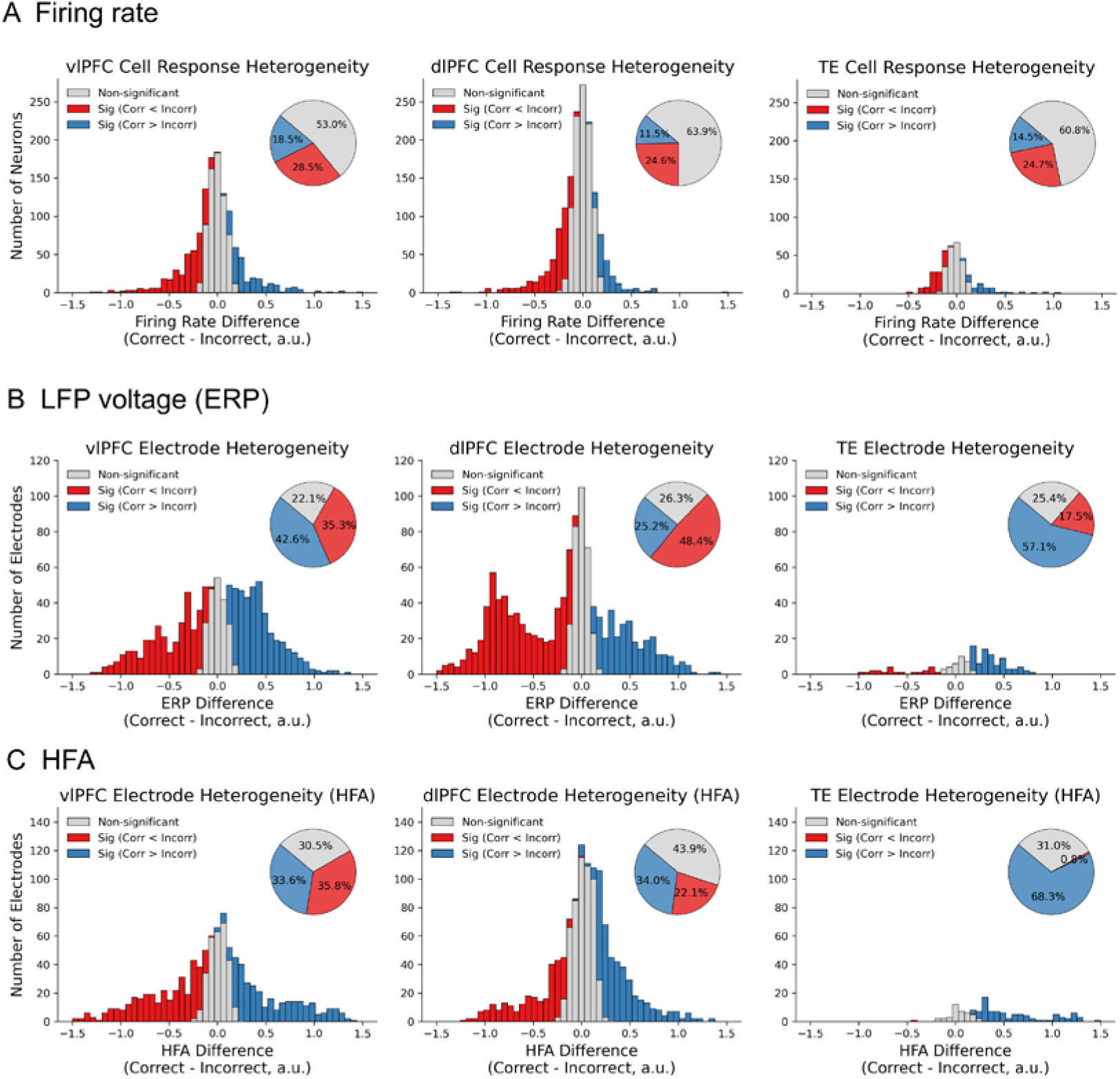
Functional heterogeneity of spiking activity, LFP voltage, and HFA. (A) Distribution of firing rate differences (Correct minus Incorrect) across all recorded neurons in vlPFC, dlPFC, and TE during the post-feedback window (200–450 ms). Stacked histograms show the number of neurons per bin, with colours indicating statistical significance (independent t-test, p < 0.05). Inset pie charts illustrate the percentage of neurons in each category. (B–C) Corresponding distributions and proportions for (B) LFP voltage and (C) high-frequency activity (HFA).

**Figure S8.**
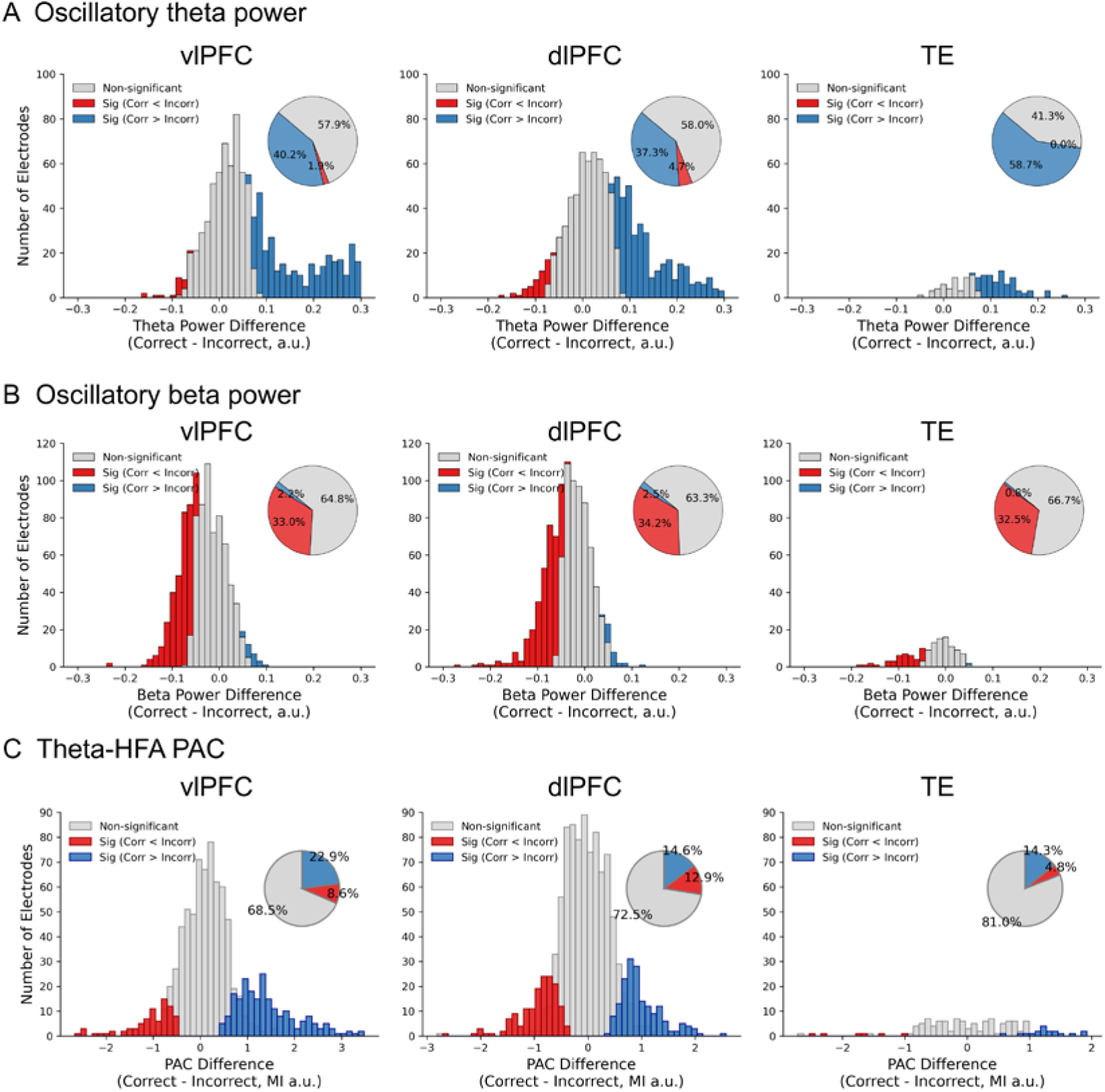
Functional heterogeneity of oscillatory power and phase-amplitude coupling. (A–B) Distributions of (A) theta (4–8 Hz) and (B) beta (20–30 Hz) power differences. (C) Distributions and proportions for theta-HFA phase-amplitude coupling (PAC). Formatting and statistical procedures are as in Figure S7.

**Figure S9.**
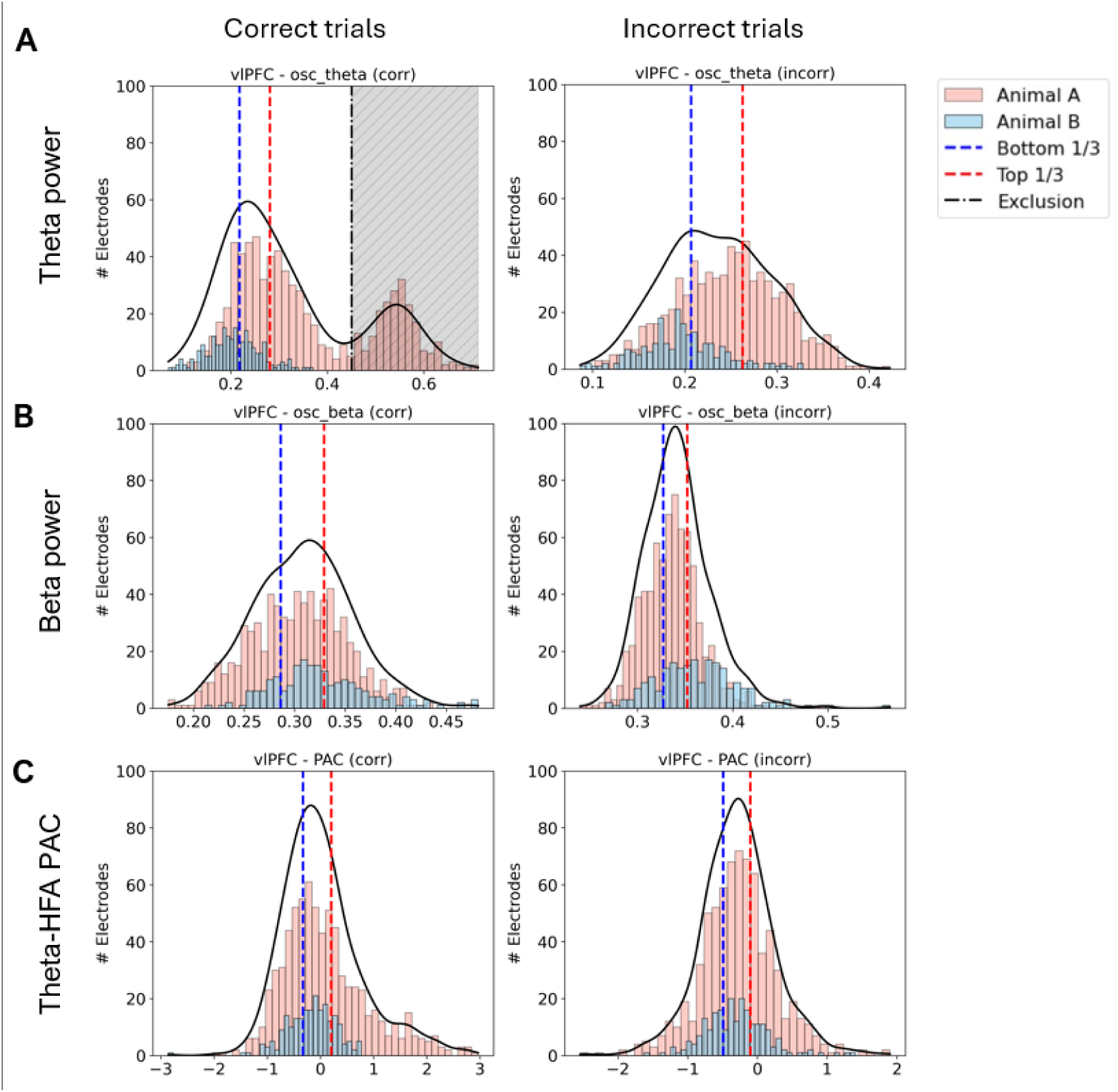
Distributions of theta, beta, and PAC values across electrodes and animals. Histograms show the distribution of (A) theta power, (B) beta power, and (C) theta-HFA phase-amplitude coupling (PAC) across all 946 vlPFC electrodes (729 from animal A, 217 from animal B). Each panel plots correct trials (left) and incorrect trials (right), based on mean activity within the 200-450 ms window following feedback onset in Cycle 1. Black lines indicate kernel-density estimates computed across both animals. Blue and red dashed lines indicate the boundaries for the bottom 1/3 and top 1/3 of electrodes, respectively, used for subsequent split analyses. Only theta power during correct feedback in animal A showed a clear bimodal distribution, with a small secondary cluster of electrodes exhibiting unusually high power. This cluster was excluded from the theta power ranking (indicated by the black dash-dotted threshold line and the grey shaded exclusion zone). Beta power and PAC showed unimodal, approximately normal distributions across both animals.

**Figure S10:**
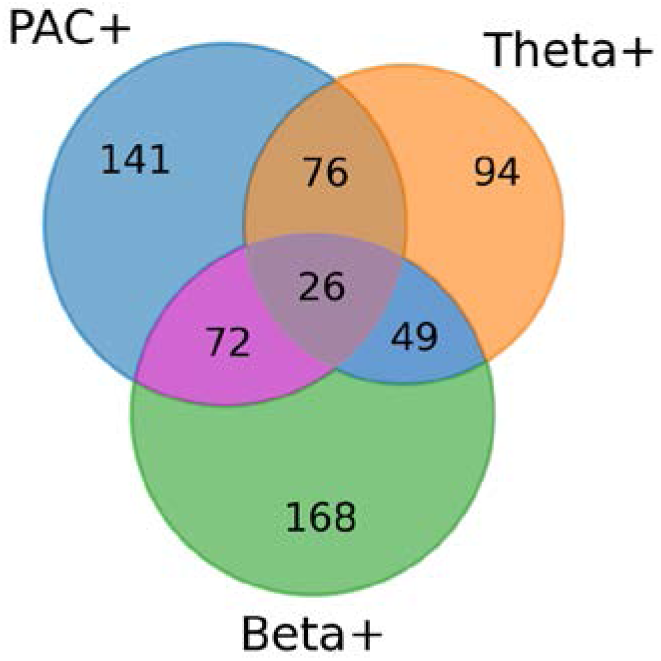
Venn diagram illustrating the modest anatomical overlap among PAC+, Theta+, and Beta+ electrode subsets in vlPFC.

**Figure S11:**
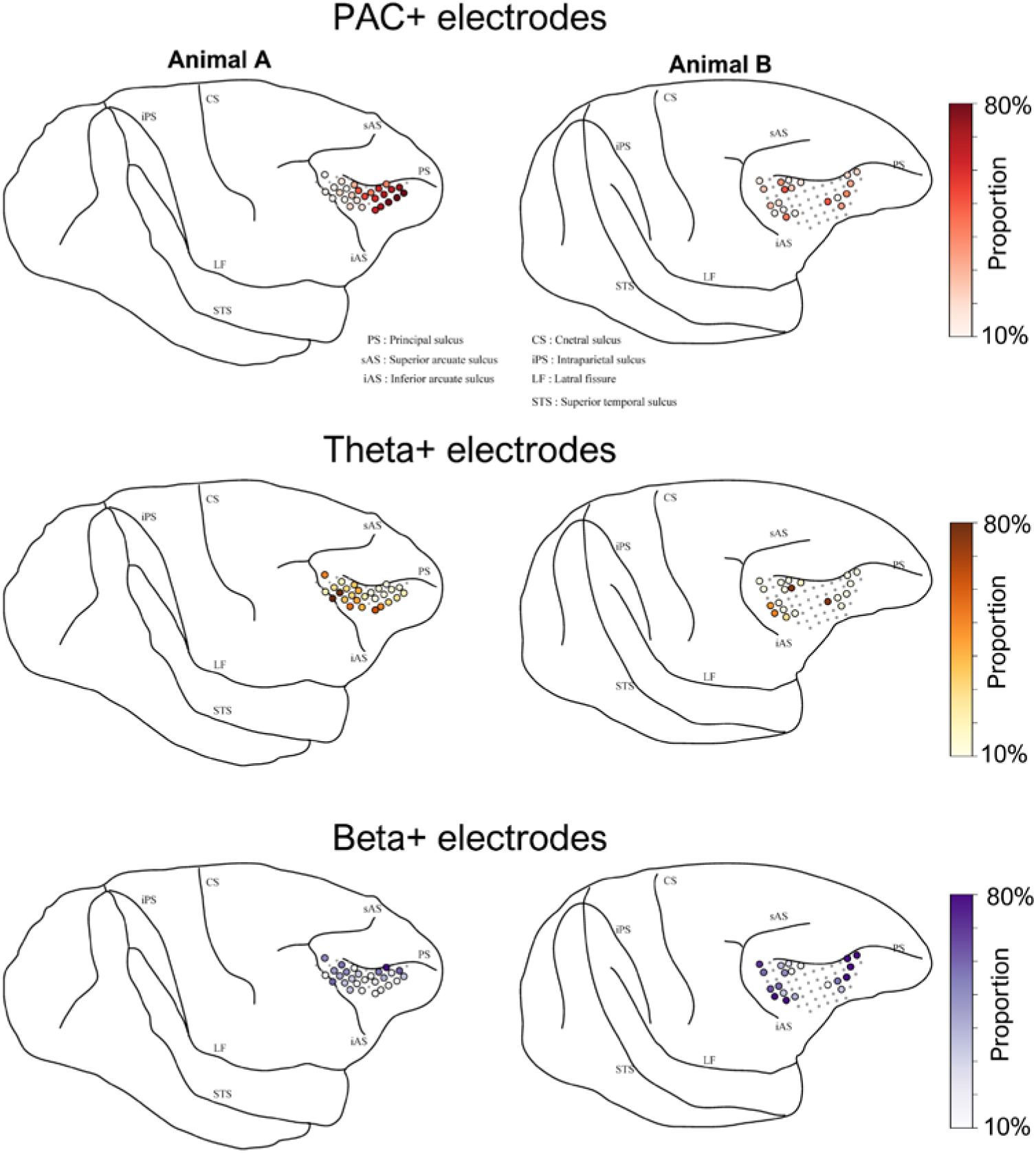
Anatomical topography of feedback-induced oscillatory signatures in the vlPFC. The spatial distribution of the electrode sites exhibiting strong PAC+, Theta+, and Beta+ responses are mapped onto the vlPFC surface for Animal A (left) and Animal B (right). The colour of each circular marker represents the proportion of different experimental sessions (corresponding to different penetration depths across days) in which a given physical electrode site ranked in the top one-third for the respective oscillatory feature. Physical electrode sites with fewer than 4 recorded sessions were excluded from this analysis.

**Supplementary Table 1.** PCA explained variance and subspace angles between Object Set 1 and Object Set 2 across neural signals and oscillatory states.

| Data Subset | Set 1<br>PC1<br>(%) | Set 1<br>PC2<br>(%) | Set 1<br>PC3<br>(%) | Set 2<br>PC1<br>(%) | Set 2<br>PC2<br>(%) | Set 2<br>PC3<br>(%) | Angle<br>1 (°) | Angle<br>2 (°) | Angle<br>3 (°) | Mean<br>Angle<br>(°) |
| --- | --- | --- | --- | --- | --- | --- | --- | --- | --- | --- |
| Spikes all | 40.0 | 33.0 | 27.0 | 38.2 | 32.5 | 29.3 | 89.9 | 88.8 | 86.5 | 88.38 |
| LFP all | 41.5 | 33.3 | 25.2 | 39.7 | 33.5 | 26.8 | 87.7 | 86.4 | 84.3 | 86.14 |
| PAC+ | 45.4 | 33.3 | 21.3 | 41.3 | 33 | 25.7 | 88.8 | 85.6 | 84.4 | 86.27 |
| PAC- | 46.7 | 28.2 | 25.1 | 37.8 | 35.5 | 26.7 | 88.6 | 77.9 | 75.6 | 80.68 |
| Theta+ | 51.1 | 26.5 | 22.4 | 43.7 | 31.4 | 24.9 | 87.2 | 84.1 | 69.6 | 80.31 |
| Theta- | 48.6 | 30.9 | 20.5 | 48 | 27 | 25 | 89.6 | 85.4 | 81.1 | 85.35 |
| Beta+ | 47.3 | 30.6 | 22.1 | 42.2 | 33.2 | 24.6 | 89 | 86.3 | 84.4 | 86.58 |
| Beta- | 46.9 | 29.1 | 24.1 | 42.2 | 31.1 | 26.7 | 90 | 86.2 | 79.5 | 85.22 |

The columns "Angle 1", "Angle 2", and "Angle 3" represent the three principal angles between the Object Set 1 and Object Set 2 subspaces within each functional data subset. The last column indicates the mean of these three angles.

